# Island Biogeography Theory-inspired predictions reveal that urban non-native plant richness is source dependent and defies classical isolation predictions

**DOI:** 10.64898/2026.08.23.746507

**Authors:** Lesibana Jan Sedibana, Yessoufou Kowiyou

## Abstract

Although cities are increasingly recognized as ecological islands, a unified framework explaining their susceptibility to alien plant invasion remains lacking.
Using the most recent and comprehensive global dataset of urban alien plants, we modelled alien richness, mimicking island biogeography theory (IBT).
Across all models, neither city size nor geographic isolation independently explained alien richness. Instead, richness was consistently associated with their interaction, supporting the central IBT prediction. However, the strength of this interaction depends on how city size was quantified, with socio-economic dimensions exhibiting stronger positive interactions with geographic isolation than physical measures of city size.
Introduction-hub identity further modified these relationships. North America was the only hub for which the interaction between city size and isolation was consistently weakened, indicating that donor regions of alien plants are not ecologically equivalent.
Simulations of simultaneous increases in city size and isolation showed that larger, more connected cities generally accumulated more alien plants despite increasing geographic distance, but the magnitude and direction of these responses are hub dependent.

Our findings inspire an extension of classical IBT to a mechanistic explanation for global variation in urban alien plant richness in this increasingly urbanized and globally connected world.

**Graphical abstract (drawn using AI):** The Urban Island Biogeography Framework (UIBF) extends classical Island Biogeography Theory to explain global patterns of alien plant richness in cities. Across six measures of urban size, neither city size nor geographic isolation independently predicted alien richness; instead, their interaction consistently explained invasion patterns. Socio-economic dimensions of urbanization (population size and GDP) strengthened this interaction more than physical city attributes, highlighting the importance of anthropogenic connectivity. Introduction-hub identity further modified these relationships, with Washington uniquely weakening the positive interaction between urban size and isolation. Together, these findings provide a mechanistic framework for predicting urban biological invasions in an increasingly connected world.

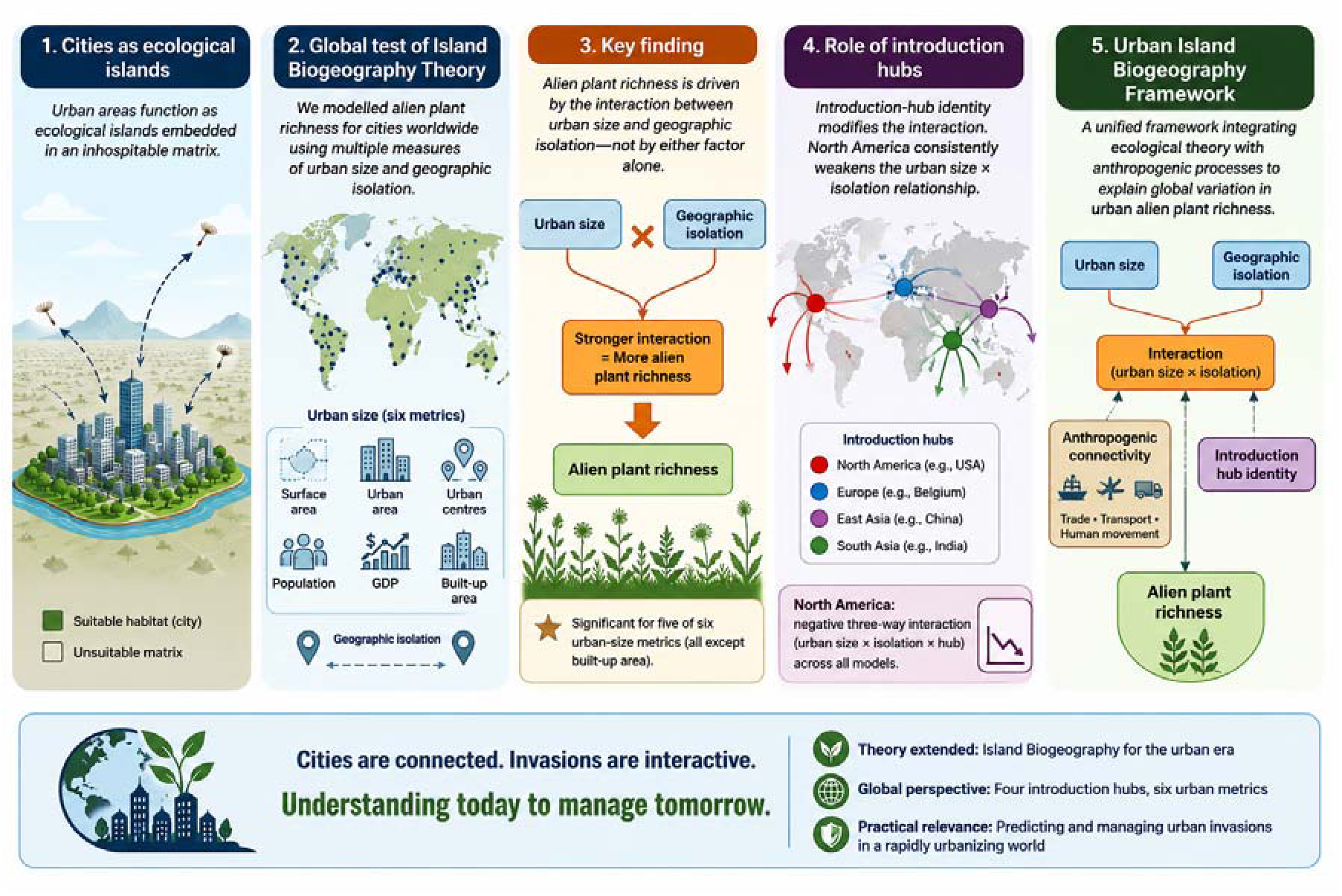

## 1. Introduction

Urbanization is among the most pervasive drivers of global environmental change in the Anthropocene. More than half of the world’s population currently resides in urban areas, and this proportion is projected to increase substantially by 2050 (United Nations, 2019). Consequently, cities are expanding rapidly, transforming natural ecosystems through land-use change, habitat fragmentation, pollution and altered hydrological regimes, thereby creating novel environmental conditions (Grimm et al., 2008; Aronson et al., 2014; Mayer et al., 2017). These anthropogenic environments facilitate the establishment of alien or non-native species that would otherwise fail to persist in surrounding natural ecosystems (Potgieter & Cadotte 2020; Richardson et al. 2025; GUBIC 2024). As urbanization accelerates worldwide, understanding the ecological processes governing biological invasions in cities has become an increasingly important challenge for ecology and conservation (Potgieter et al. 2024).

Cities are now acknowledged as global hotspots of non-native plant diversity (Aronson et al. 2015; Cadotte et al. 2017; Potgieter et al. 2020; Potgieter, Li, et al. 2024; Li et al. 2025). Beyond supporting high numbers of alien species, they also function as important sources from which non-native plants spread into adjacent natural and semi-natural ecosystems (McLean et al. 2017; Tartaglia et al. 2018; Potgieter, ter Huurne, and Richardson 2024). Although urban biological invasions have been extensively investigated in Europe (e.g., Sukopp and Werner 1983; Kowarik 1990; Sukopp et al. 1990; Pyšek 1993, 1998; Sukopp 2002; Celesti-Grapow et al. 2006; Lososová et al. 2012), our understanding remains comparatively limited outside Europe, including North America (Aronson et al. 2007, 2015), Africa (Gaertner et al. 2017), New Zealand (Ricotta et al. 2017) and Latin America (Figueroa et al. 2018). Consequently, despite considerable progress, a general understanding of the processes governing urban invasions across biogeographic regions is still lacking (but see Potgieter et al. 2024).

The ecological and socioeconomic consequences of urban biological invasions further emphasize the need to understand why some cities are more susceptible to invasion than others. Non-native species alter ecosystem structure and functioning (Pyšek et al. 2020) while increasing ecosystem disservices in urban environments (Potgieter et al. 2017; Vaz et al. 2017; Milanović et al. 2020). These include allelopathic effects, allergenic pollen production, soil erosion, habitat modification and damage to urban infrastructure (Vaz et al. 2017; Milanović et al. 2020). As urban areas continue to expand globally, these impacts are expected to intensify (Perrings et al. 2010; Roy et al. 2023; Potgieter et al. 2024). Consequently, substantial efforts have been devoted to preventing and managing urban biological invasions (e.g., Gaertner et al., 2016; Potgieter et al., 2018; Potgieter & Cadotte, 2020; Potgieter et al., 2022, 2024; Gildenhuys et al., 2025). However, effective management depends not only on reactive interventions but also on understanding the ecological mechanisms that determine why some cities accumulate substantially more non-native species than others. Although recent global syntheses have identified several correlates of urban invasibility, including geographic connectivity, city size, socioeconomic conditions and biogeography (Potgieter et al., 2024), these determinants remain largely descriptive. Urban invasion ecology and policy makers would benefit more from a research driven unifying theoretical framework capable of integrating these multiple drivers into a coherent explanation of global variation in urban alien species richness.

Island Biogeography Theory (IBT; MacArthur and Wilson 1967) provides an attractive conceptual foundation for addressing this challenge. IBT predicts that species richness reflects the balance between immigration and extinction, such that larger islands located closer to the mainland support more species than smaller and more isolated islands. This prediction has been repeatedly supported across a wide range of insular systems, including oceanic islands, ponds and fragmented habitats (Rosenzweig 1995; Whittaker & Fernández-Palacios, 2007; Whittaker et al., 2017). At the same time, the theory has evolved through a series of conceptual refinements that recognized the importance of additional ecological and evolutionary processes influencing species richness (e.g. Brown & Lomolino, 2000; Lomolino, 2000; Heaney, 2007). More recently, there have been calls to extend the theory by explicitly incorporating anthropogenic processes that may fundamentally modify immigration dynamics and potentially override the effects of geographic isolation (Liu et al. 2023). Such developments are particularly relevant for urban ecosystems, where species introductions are overwhelmingly mediated by human activities rather than natural dispersal (Potgieter et al. 2024; Richardson et al. 2025).

Cities, indeed, share several characteristics with islands. They represent discrete habitat patches embedded within a surrounding matrix that is often less suitable for many non-native species, and they differ markedly in size, connectivity and environmental conditions. However, unlike oceanic islands, immigration into cities is largely human-aided, including trade, transport, horticulture and economic exchange. Consequently, geographic isolation may not necessarily reduce immigration as predicted by classical IBT. Instead, large and economically important cities may continue to receive substantial numbers of introduced alien species, despite being geographically distant from their primary sources of introduction. This possibility raises a fundamental question: does Island Biogeography Theory adequately explain urban biological invasions, or do human-mediated dispersal processes alter its central predictions?

Addressing this question also requires recognizing that species introductions are not spatially uniform. Cities receive non-native species disproportionately from different regions of the world through distinct historical, commercial and cultural pathways (GUBIC 2025; Richardson et al. 2025). These regions, referred to, here, as *introduction hubs*, differ in propagule pressure, donor species pools, introduction histories and climatic affinities, suggesting that the relationship between urban size, geographic isolation and alien plant richness may depend on the origin of introduced species. Globally, temperate and tropical Asia constitute by far the largest introduction hub of alien plants into cities, followed by Europe (Richardson et al. 2025). This uneven contribution of donor regions raises the possibility that introduction-hub identity might modify the predictions of Island Biogeography Theory in urban ecosystems. Consequently, the interaction between city size and geographic isolation may not be universal but instead vary among introduction hubs. Incorporating source-dependent immigration into an island biogeographic framework therefore offers a mechanistic approach for understanding why cities with similar characteristics may differ substantially in their non-native species richness.

Historically, testing such a framework has been hindered by the lack of harmonized global datasets. Previous studies have typically focused on individual cities, limited geographic regions or specific taxonomic groups (Vaz et al. 2018), while collection methods, taxonomic standards and sampling efforts have differed considerably among studies (GUBIC 2025). Consequently, attempts to identify global drivers of urban invasions have remained challenging (see Gaertner et al., 2016; Potgieter & Cadotte, 2020; Cadotte et al. 2007). The recently developed Global Urban Biotic Invasions Compendium (GUBIC 2025) overcomes many of these limitations by providing a harmonized global repository of non-native species records across cities worldwide, creating an unprecedented opportunity to evaluate broad-scale hypotheses in urban invasion ecology.

Using the GUBIC dataset, we investigate whether Island Biogeography Theory provides a predictive framework for explaining global patterns of urban alien plant richness. Specifically, we test whether (1) Island Biogeography Theory explains variation in alien plant richness among cities, (2) alternative dimensions of city size, including physical, demographic and socioeconomic measures, differ in their influence on urban alien plant richness, (3) increasing city size modifies the effects of geographic isolation on alien plant richness, and (4) the relationships among city size, geographic isolation and alien plant richness depend on the introduction hub from which species originate. By integrating classical biogeographic theory with human-mediated pathways of species introduction, our study evaluates whether one of ecology’s most influential theories can explain biological invasions in cities or whether urban ecosystems require a modified biogeographic framework.

## 2. Materials and Methods

### 2.1 Study system

This study was conducted at a global scale (Fig. 1) using data from 553 urban centres distributed across 61 countries on all continents except Antarctica (GUBIC 2025). In the GUBIC dataset, urban centres were defined following the Global Human Settlement Layer (GHSL) as contiguous 1-km² grid cells with a population density of at least 1,500 inhabitants km⁻² or >50% built-up area, and a minimum population of 50,000 inhabitants (Li et al. 2025).

**Figure 1.**
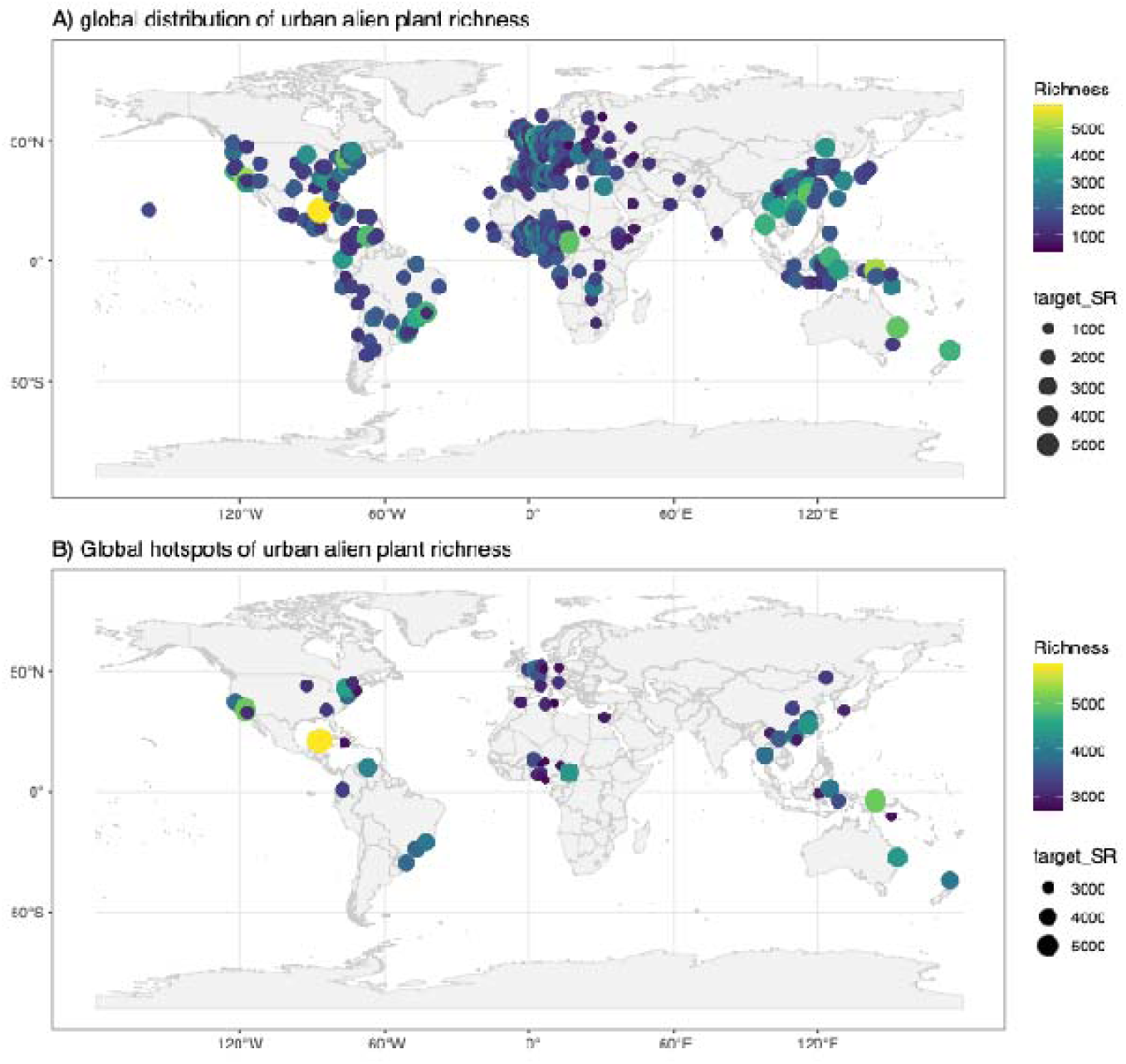
Global distribution of urban alien plant richness illustrating. (A) urban plant species richness and (B) hotspots of urban alien plant richness represented by cities within the upper 10% of the alien richness distribution, highlighting the major global centres of urban plant invasion. Point colour and size are proportional to the number of recorded alien plant species, illustrating substantial spatial variation in urban invasion richness worldwide.

### 2.2. Conceptual source-target networks: selection of representative introduction hubs

To evaluate whether Island Biogeography Theory (IBT) explains global patterns of urban alien plant richness, we developed a conceptual source–target network in which four globally connected metropolitan areas served as representative introduction hubs (Fig. 2). These hubs were selected to represent the principal global donor regions of alien plants identified by Richardson et al. (2025), who showed that temperate and tropical Asia constitute by far the largest source region of alien plants worldwide, followed by Europe. Accordingly, Beijing (China) and New Delhi (India) were selected to represent two major Asian introduction hubs, Brussels (Belgium) represented Europe, and Washington (USA) represented the Americas. Washington was chosen because North America contributes substantially more alien plant species globally than South America, which is a comparatively minor donor region (Richardson et al. 2025).

**Figure 2.**
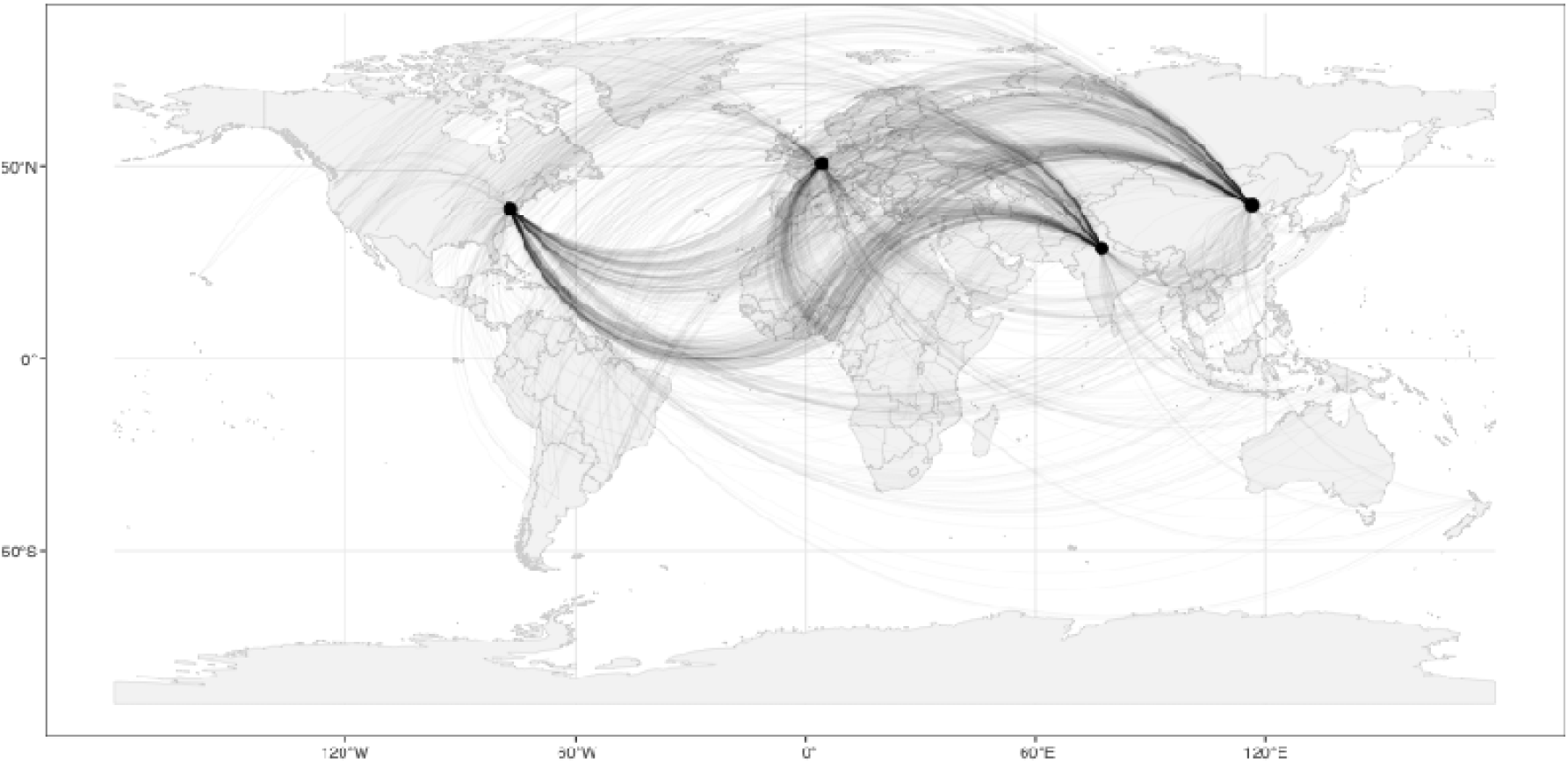
Conceptual source–target network used to quantify geographic isolation in the Urban Island Biogeography Framework. Four globally connected metropolitan areas (Beijing, Brussels, New Delhi and Washington) were selected as representative introduction hubs of alien plants based on their major contributions to global plant exchange (Richardson et al. 2025). Geographic isolation was calculated as the great-circle distance between each target city and each introduction hub, with each curve representing one source–target relationship used in the analyses. All curves are plotted with identical line width and transparency; regions where curves appear thicker represent the overlap of multiple source-target connections rather than differences in connection strength or invasion intensity.

These cities were not assumed to be the exclusive sources of the alien plants recorded worldwide. Rather, they were used as representative proxies for major donor regions because they are globally connected metropolitan centres with long histories of international trade, transportation, horticultural exchange and plant introductions. Their selection also reflects the importance of biogeographic context in shaping urban biological invasions (Potgieter et al. 2024). Together, these cities capture distinct historical, economic and biogeographic pathways through which alien plants have been redistributed globally.

Within the conceptual source–target network (Fig. 2), the four metropolitan centres represented the mainland, whereas all remaining cities were treated as urban islands. Geographic isolation was therefore quantified as the great-circle distance between each target city and each representative introduction hub, allowing us to evaluate whether the relationship between urban size and alien plant richness varies according to both geographic isolation and the identity of the donor region.

### 2.3 Data collection

#### 2.3.1 Alien plant richness

Data on non-native vascular plants were retrieved from the Global Urban Biological Invasion Compendium (GUBIC; https://doi.org/10.5281/zenodo.14559926; accessed June 2026), currently the most comprehensive global database of alien plant diversity in urban ecosystems (Li et al. 2025; Richardson et al. 2025). GUBIC integrates information from multiple complementary data sources to provide a harmonized global inventory of urban alien plants. First, members of the GUBIC consortium contributed urban species records through a centralized SharePoint repository. Second, published studies in English, Spanish and Portuguese were systematically reviewed, and additional datasets archived in the Dryad repository were incorporated. Third, records were extracted from the Urban Biodiversity Research Coordination Network (UrbioNet). Finally, occurrence records in the database were obtained from the Global Biodiversity Information Facility (GBIF; Li et al. 2025). The integration of these complementary data sources produced the most comprehensive global compilation of urban non-native vascular plants currently available.

The final dataset comprises 8,140 alien vascular plant species belonging to 253 families recorded across 553 urban centres in 61 countries spanning all continents except Antarctica. Urban boundaries followed the urban-centre delineation of the Global Human Settlement Layer (GHSL; Li et al. 2025).

#### 2.3.2 City characteristics

City characteristics were obtained from the Global Human Settlement Layer (GHSL) (Pesaresi et al., 2019; https://ghsl.jrc.ec.europa.eu/ucdb2018Overview.php). We restricted our data collection on cities to the urban centres included in GUBIC for which alien plant richness data were available.

##### City size

Because urban size is multidimensional, we quantified it using several complementary descriptors available from GHSL and incorporated into GUBIC. These included: city surface area (km²; year 2015); number of urban centres (year 2000); total urban area (km²; year 2000); total built-up area (km²; year 2015); population size (year 2015); and Gross Domestic Product (GDP; year 2015). Together, these variables capture the physical, demographic and socioeconomic sizes of all cities.

##### Geographic isolation

Within the IBT framework, geographic isolation was quantified as the great-circle distance between each introduction hub and every target city. Distances were calculated between the centroids of GHSL urban centres using the st_distance function implemented in the sf package (Pebesma, 2018) in R (Version 4.5.1; R Core Team, 2025). Separate distance matrices were generated for each of the four introduction hubs, allowing the influence of geographic isolation to be evaluated independently for each putative source region.

### 2.4 Data analysis

All data analyzed are available in FigShare (see Sedibana et al. 2026). All statistical analyses were conducted in R (Version 4.5.1; R Core Team, 2025). Continuous predictor variables, including geographic isolation, city surface area, urban area, total built-up area, population size and gross domestic product (GDP), were standardized (mean = 0, SD = 1) prior to modelling to facilitate model convergence and comparison of effect sizes. The number of urban centres was retained on its original scale because it represents a discrete count variable.

To evaluate whether Island Biogeography Theory (IBT) explains global variation in urban alien plant richness, we modelled non-native plant species richness as a function of city size, geographic isolation and introduction hub using negative binomial generalized linear mixed models fitted with the glmmTMB package (our response variable, species richness, is a count data with overdispersion). Species richness was modelled assuming Gaussian errors, and country was included as a random intercept to account for the non-independence of cities within countries.

Because urbanization encompasses multiple dimensions, we evaluated six complementary measures of city size in separate models: (1) city surface area, (2) urban area, (3) number of urban centres, (4) total built-up area, (5) human population size and (6) gross domestic product (GDP). Geographic isolation was quantified as the great-circle distance between each target city and one of four representative introduction hubs (Beijing, Brussels, New Delhi and Washington).

The general model structure was:

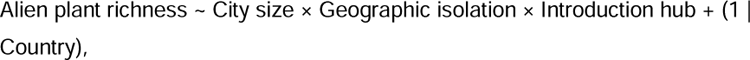

where *alien plant richness* is the number of non-native plant species recorded in a city; *city size* was represented, in separate models, by (1) city surface area, (2) urban area, (3) number of urban centres, (4) total built-up area, (5) population size, or (6) gross domestic product (GDP); *geographic isolation* is the great-circle distance between the target city and the assigned introduction hub; *introduction hub* comprises the four representative source cities (Beijing, Brussels, New Delhi and Washington); and *country* was included as a random intercept.

The modelling strategy was designed to address the four objectives of the study. First, the main effects of city size and geographic isolation were used to evaluate whether patterns of urban alien plant richness conform to the classical predictions of IBT. Second, fitting separate models for alternative measures of city size allowed us to determine whether different dimensions of urbanization differ in their ability to explain alien plant richness. Third, the interaction between city size and geographic isolation tested whether urbanization modifies the effect of isolation on species richness, thereby assessing whether urban growth mitigates or amplifies the isolation effect predicted by IBT. Finally, the three-way interaction among city size, geographic isolation and introduction hub tested whether these relationships depend on the geographic origin of introduced species, thereby evaluating whether the applicability of IBT varies among source regions.

To facilitate biological interpretation of significant interactions, marginal predictions were generated from each fitted model across the observed gradient of geographic isolation while holding each city-size variable at representative values spanning its observed range. Predictions were generated separately for each introduction hub, and standardized geographic distances were back-transformed to kilometres for graphical presentation.

To further examine how the influence of urbanization varies along the isolation gradient, conditional marginal effects of each city-size variable were estimated across the observed range of geographic isolation using the marginaleffects package. These analyses quantified how the effect of increasing city size on alien plant richness changes with increasing geographic isolation for each introduction hub.

To compare the relative importance of different urbanization metrics, regression coefficients describing the main effects of city size and geographic isolation, the two-way interaction between city size and geographic isolation, and the three-way interaction involving introduction hub were extracted from each fitted model and summarized graphically. This synthesis enabled direct comparison of the magnitude, direction and statistical significance of the processes underlying urban alien plant richness across alternative measures of city size.

Finally, because standardized regression coefficients are difficult to interpret ecologically, we quantified the biological consequences of the interaction effects using model-based predictions. For each fitted model, alien plant richness was first predicted for a reference city with all continuous predictors fixed at their mean values. Predictions were then recalculated after simultaneously increasing geographic isolation by 1000 km and one measure of city size by an ecologically meaningful increment, namely 100 km² for city surface area, urban area and built-up area, one additional urban centre, one million inhabitants, or US$10 billion GDP. The difference between the two predictions was interpreted as the expected change in alien plant richness associated with simultaneous increases in urbanization and geographic isolation. These calculations were performed separately for each introduction hub to quantify how the ecological consequences of urban expansion differ among source regions.

## 3. Results

### 3.1 Urban size and geographic isolation jointly explain global patterns of alien plant richness

Across all six negative binomial mixed models, neither urban size nor geographic isolation independently explained variation in alien plant richness (all *P* > 0.05; Figure 3). Instead, support consistently emerged for an interaction between urban size and geographic isolation, indicating that the influence of city size depended on the degree of isolation from the introduction hub. The interaction between urban size and isolation was strongest for socio-economic measures of city size. Significant positive interactions were detected when urban size was represented by population size (β = 0.052, *P* = 0.027), gross domestic product (GDP) (β = 0.083, *P* = 0.032) and the number of urban centres (β = 0.077, *P* = 0.038). Similar but marginally significant interactions were observed for city surface area (β = 0.061, *P* = 0.076) and urban area (β = 0.065, *P*= 0.060), whereas no evidence of an interaction was detected for built-up area (β = 0.062, *P* = 0.114; Figure 3). These results indicate that larger cities generally accumulated more alien plant species with increasing geographic isolation, but that the strength of this relationship depended on how urban size was quantified.

**Figure 3.**
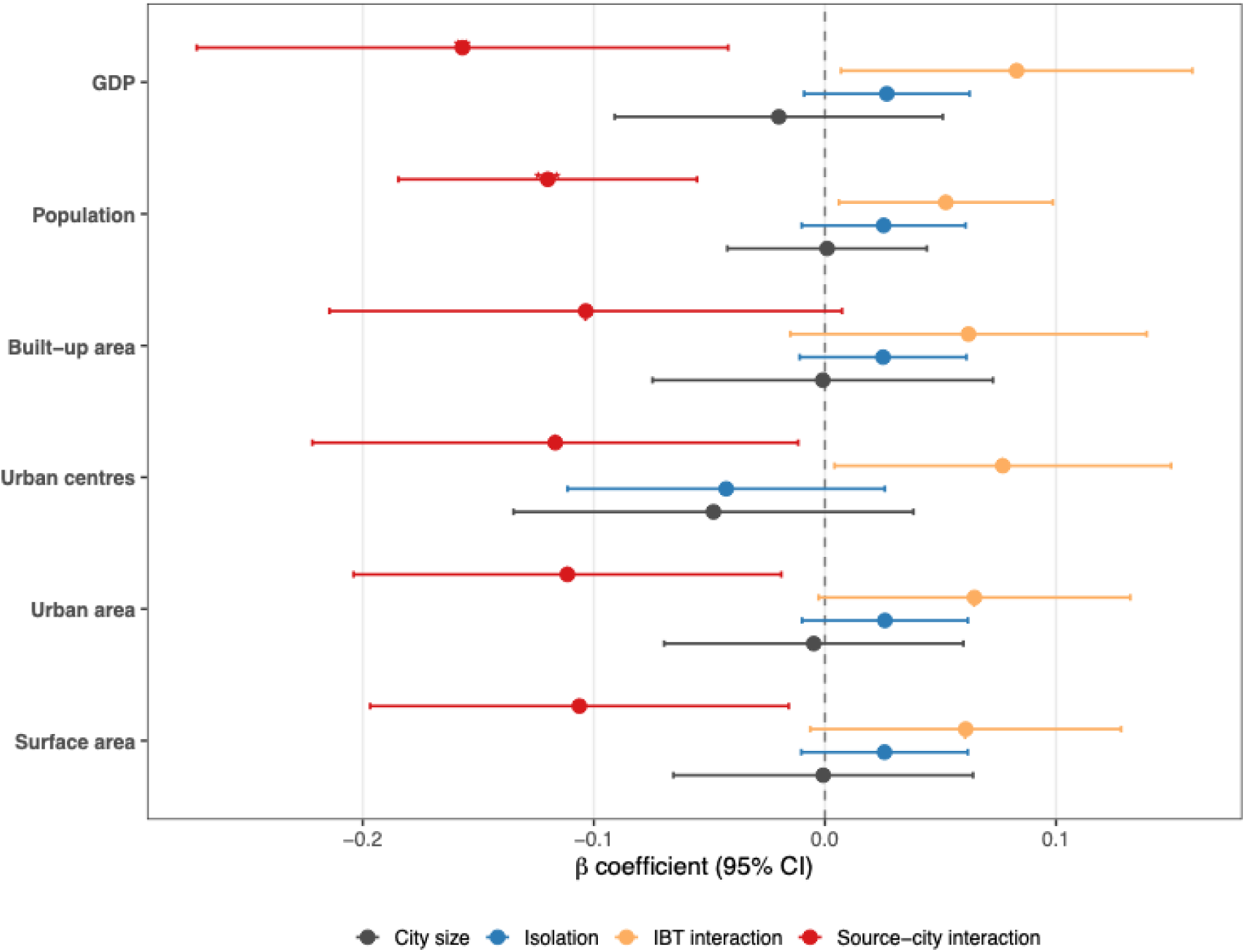
Consistency of Island Biogeography Theory (IBT) effects across alternative measures of urban size. Forest plot showing standardized regression coefficients (β ± 95% confidence intervals) from six negative binomial generalized linear mixed models in which urban size was represented by city surface area, urban area, number of urban centres, built-up area, population size, and gross domestic product (GDP). Four effects are compared across all models: the direct effect of city size (black), the direct effect of geographic isolation (blue), the city size × isolation interaction predicted by classical Island Biogeography Theory (orange), and the three-way interaction between city size, geographic isolation and introduction-hub identity (Washington relative to Beijing; red). The vertical dashed line indicates no effect (β = 0). Positive two-way interaction coefficients indicate that the influence of city size on alien plant richness increases with geographic isolation, whereas negative three-way interaction coefficients indicate that this relationship is significantly weakened when Washington is considered the introduction hub.

### 3.2. Introduction-hub identity modifies the relationship between urban size and isolation

The interaction between urban size and geographic isolation was further modified by the identity of the introduction hub. Across all six models, Washington was the only introduction hub exhibiting consistently negative three-way interactions, indicating that the positive association between urban size and isolation weakened when North America was considered the donor region. This moderating effect was significant when urban size was represented by physical parameters of cities (Figure 4): city surface area (β = −0.106, *P* = 0.022), urban area (β = −0.111, *P* = 0.018), and number of urban centres (β = −0.117, *P* = 0.030).

**Figure 4.**
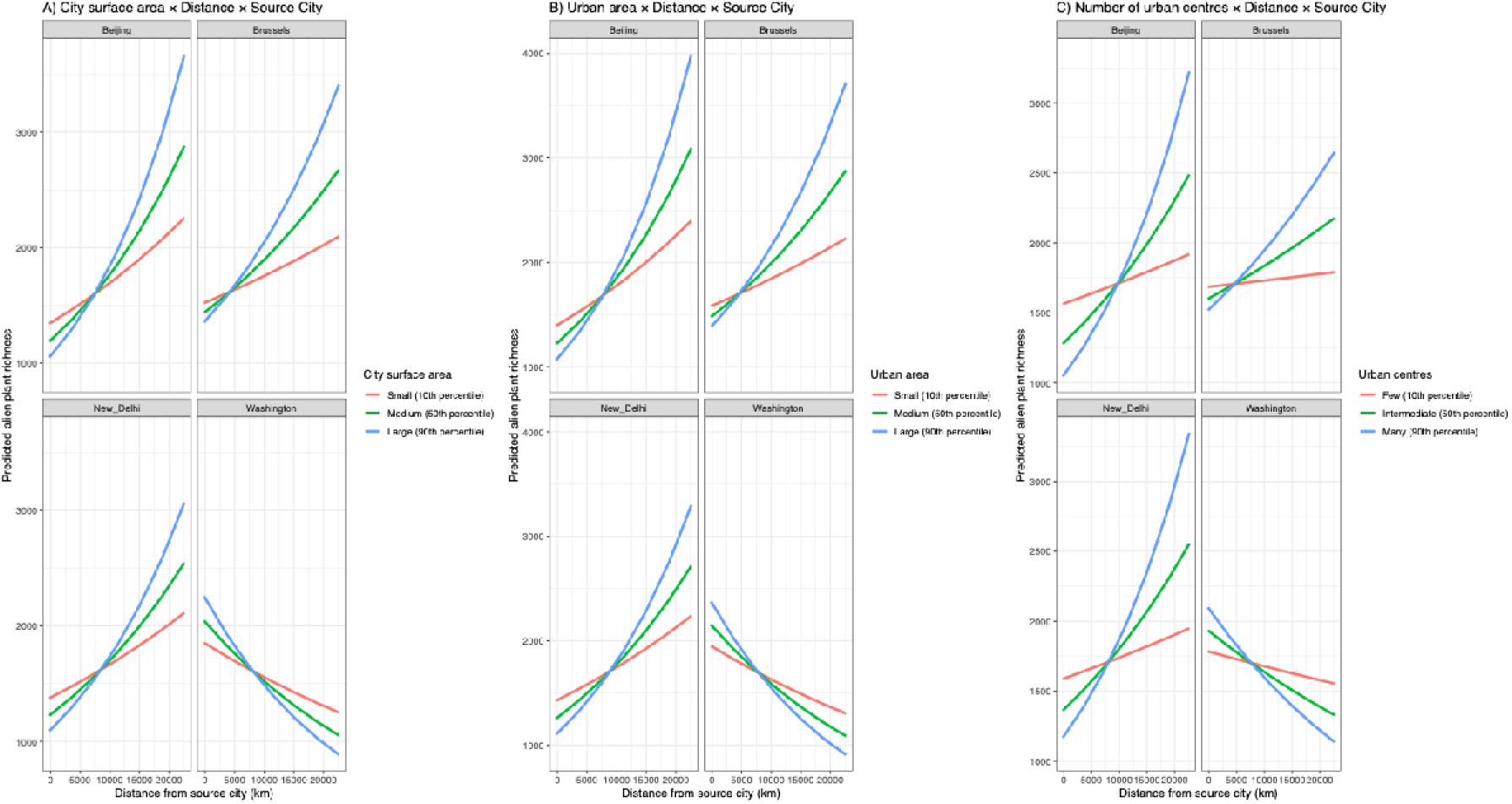
Effects of physical variables of cities, geographic isolation, and source city on alien plant species richness across the six generalized linear mixed models. These physical variables include city surface area, urban area and number of urban centres.

Similar patterns were found when socio-economic parameters were used to approximate city size (Figure 5): population size (β = −0.120, *P* < 0.001) and GDP (β = −0.157, *P* = 0.007). Built-up area showed the same negative tendency but did not reach statistical significance (β = −0.103, *P* = 0.067). No significant three-way interactions were detected for Beijing, Brussels or New Delhi (Table 1). These findings indicate that the effect of geographic isolation on alien plant richness is source-region dependent rather than universal.

**Figure 5.**
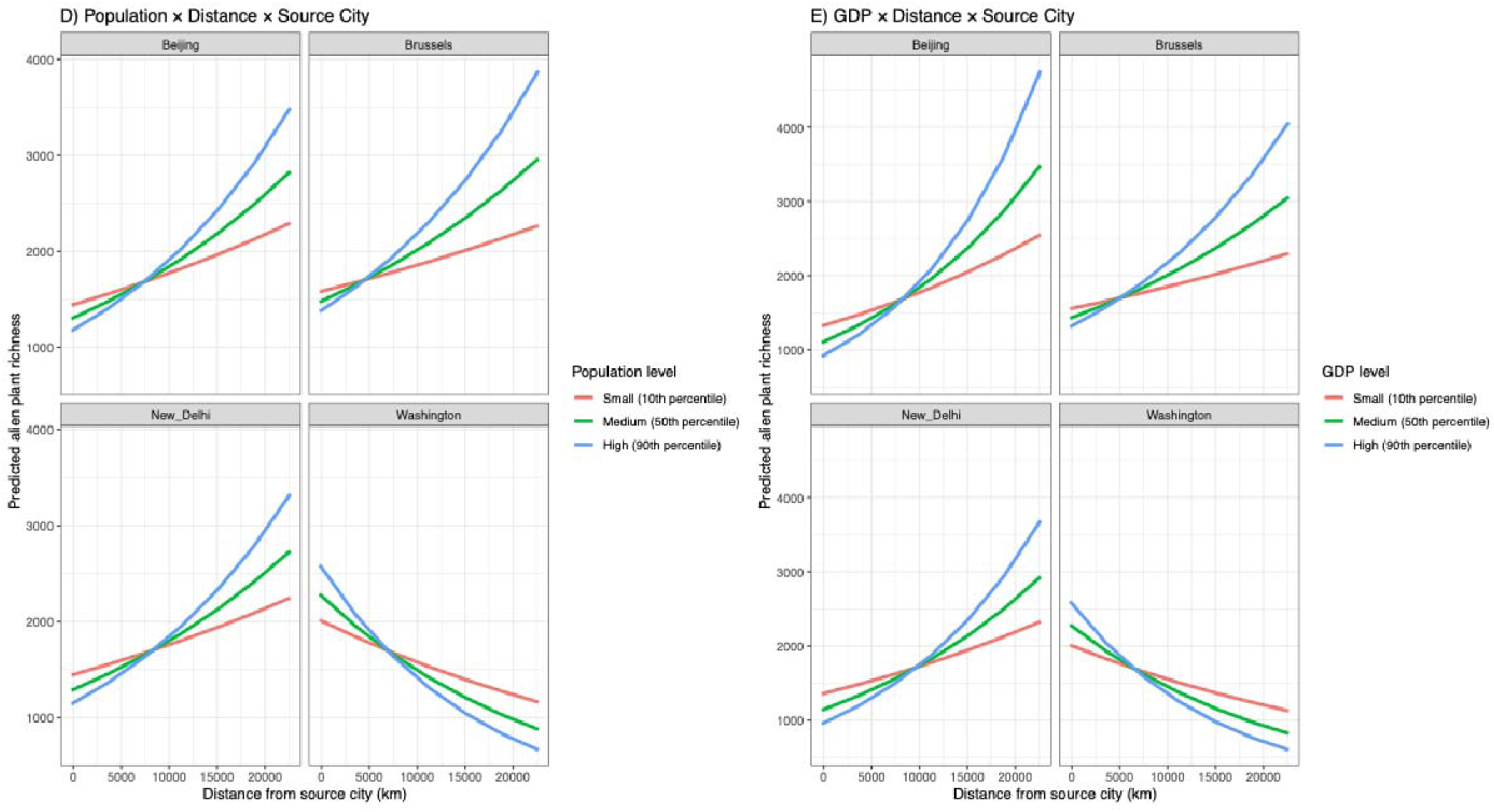
Effects of socio-economic variables of cities, geographic isolation, and source city on alien plant species richness across the six generalized linear mixed models. These physical variables include population size and GDP.

### 3.2 Predicted effects of simultaneous increases in urban size and geographic isolation

To illustrate the ecological implications of the fitted models, we simulated simultaneous increases in urban size and geographic isolation by increasing each urban-size metric while increasing the distance between introduction hub and recipient city by 1000 km. The predicted responses differed markedly among introduction hubs and measures of urban size (Figure 6).

**Figure 6.**
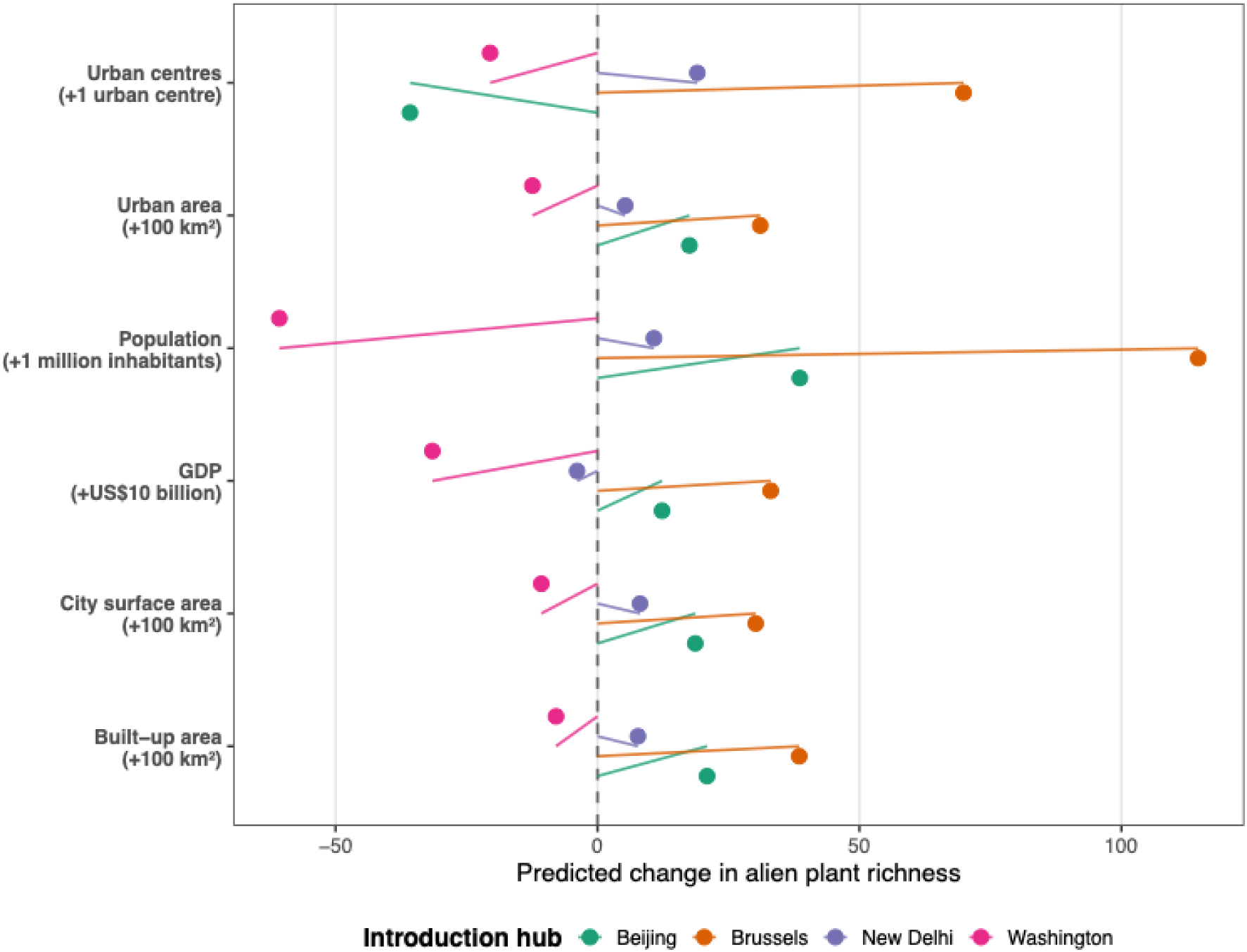
Forest plot showing the predicted change in alien plant richness associated with simultaneous increases in urban size and geographic isolation (1000 km) for each introduction hub. Urban-size increments were 100 km² for city surface area, urban area and built-up area, one additional urban centre, one million inhabitants for population size, and US$10 billion for GDP. Positive values indicate increases in predicted alien plant richness relative to the reference condition, whereas negative values indicate decreases. The contrasting responses among introduction hubs demonstrate that the effects of urban expansion on alien plant richness depend strongly on source-region identity.

Across all physical measures of urban size (city surface area, urban area and built-up area), Brussels consistently exhibited the largest increases in predicted alien plant richness, with gains ranging from 30.2 to 38.5 species, whereas Beijing showed moderate increases (17.5-20.9 species) and New Delhi exhibited relatively small increases (5.3-8.1 species). In contrast, Washington consistently showed declines in predicted richness despite the increase in urban size, with losses ranging from 7.9 to 12.4 species. Predictions based on the number of urban centres revealed greater variation among introduction hubs. Brussels again showed the largest increase (+69.9 species), followed by New Delhi (+19.1 species), whereas Beijing (−35.8 species) and Washington (−20.5 species) both exhibited declines in predicted richness.

The strongest contrasts were observed for the socio-economic measures of urban size. Increasing city population by one million inhabitants while simultaneously increasing geographic isolation by 1000 km resulted in a predicted increase of 114.7 species for Brussels, 38.6 species for Beijing and 10.8 species for New Delhi. In contrast, Washington showed the largest decline across all simulations (−60.7 species). Similarly, increasing GDP by US$10 billion produced moderate increases for Brussels (+33.0 species) and Beijing (+12.3 species), but resulted in declines for New Delhi (−3.9 species) and Washington (−31.5 species).

Overall, based on all our findings, we proposed a framework to explain alien invasion in urban ecosystems (Figure 7). In this framework, urban size is represented by multiple dimensions (surface area, urban area, number of urban centres, population size, GDP, and built-up area), with significant support for the interaction across all metrics except built-up area. Anthropogenic connectivity, including trade, transportation, and human movement, increases both urban growth and the exchange of propagules among cities, thereby modifying the influence of geographic isolation on colonization success. The identity of the introduction hub further alters this relationship, with Washington exhibiting a consistently negative three-way interaction between urban size and isolation across all models, indicating that source-region identity influences how geographic isolation affects invasion dynamics. Dashed arrows represent weak or non-significant direct effects, whereas solid arrows indicate supported relationships. Together, these findings extend classical Island Biogeography Theory by explicitly incorporating anthropogenic connectivity and source-region effects into a unified framework for understanding and predicting alien plant richness in cities worldwide.

**Figure 7.**
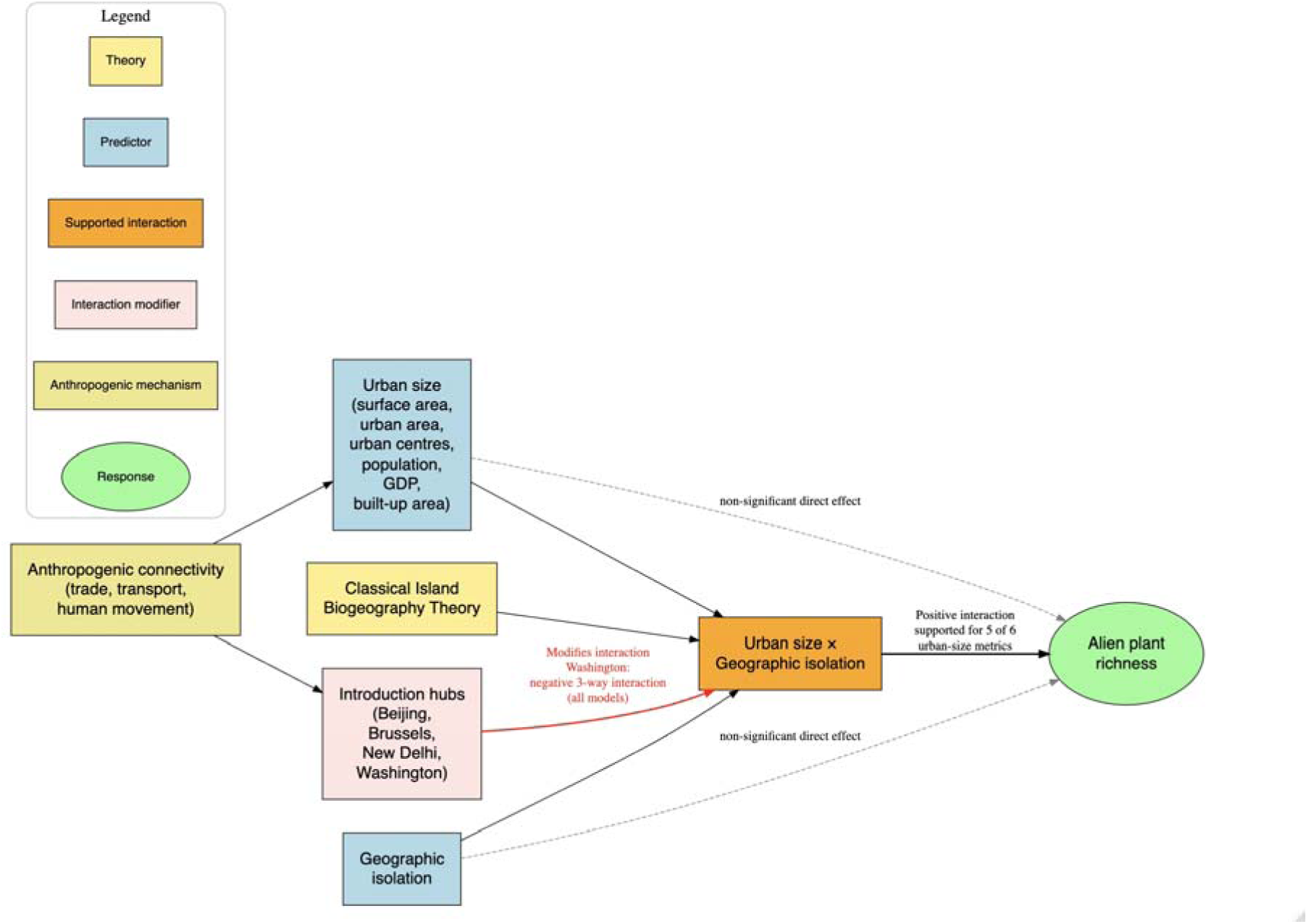
Proposed Urban Island Biogeography Framework (UIBF) integrating classical Island Biogeography Theory with anthropogenic processes driving biological invasions in cities.

## Discussion

### Urban Island Biogeography Theory extends classical Island Biogeography Theory to cities

The central objective of this study was to evaluate whether the core predictions of Island Biogeography Theory (IBT) can explain global patterns of alien plant richness in cities. Our results provide qualified support for this hypothesis. Across six independent models representing different dimensions of urban size, neither urban size nor geographic isolation consistently explained alien plant richness when considered independently. Instead, alien richness was repeatedly explained by the interaction between urban size and geographic isolation, indicating that these processes operate jointly rather than independently. This pattern closely reflects the central premise of classical IBT, namely that species richness emerges from the combined effects of habitat area and isolation rather than from either process alone (MacArthur & Wilson 1967). Our findings therefore suggest that cities function as ecological islands, while simultaneously highlighting that urban systems require an extension of classical IBT that explicitly incorporates anthropogenic processes governing species movement (Liu et al. 2023).

The direction of the interaction, however, differed from the expectation of classical IBT. MacArthur and Wilson (1967) predicted that increasing isolation reduces immigration and consequently species richness. In contrast, we found that geographic isolation alone had little explanatory power, whereas its interaction with urban size was consistently positive across most models. This result indicates that geographic distance no longer functions as an absolute barrier to plant dispersal in highly connected urban systems. Instead, the influence of isolation appears contingent on city characteristics, particularly those associated with human activities (Potgieter et al. 2024). This implies that large cities receive disproportionately more propagules likely through trade, transportation and human mobility, thereby reducing the constraining effect of geographic distance (O’Malia et al. 2018; Potgieter et al. 2024). Also, urbanization creates greater habitat heterogeneity within larger cities, potentially further enhancing establishment opportunities for introduced species as predicted in the habitat amount hypothesis (Callaghan et al. 2019). Overall, geographic isolation remains relevant, but its ecological effects are increasingly mediated by anthropogenic connectivity rather than by distance alone.

An important outcome of our analyses is that support for the urban size-isolation interaction was remarkably consistent across all types of metrics quantifying city size. Whether city size was represented by city surface area, urban area, number of urban centres, population size or GDP, the interaction was always positive, although its statistical strength varied among metrics. Only built-up area failed to show evidence of a significant interaction. This consistency suggests that the observed relationship is not an artefact of a particular definition of city size but reflects a general ecological process governing alien plant accumulation across cities.

### Socio-economic dimensions of city size better explain alien plant richness

A major contribution of this study is demonstrating that urban size is multidimensional and that different dimensions contribute unequally to biological invasions. Classical applications of IBT almost exclusively consider island area as a surrogate for habitat availability. Cities, however, are simultaneously spatial, demographic and socio-economic systems (Potgieter & Cadotte 2020; Potgieter et al. 2024). Consequently, a single measure of urban size is unlikely to capture all processes influencing species accumulation.

Consistent with previous urban ecology studies, physical measures of city size, including surface area and urban extent, showed positive interactions with geographic isolation, supporting their long-recognized importance in explaining urban biodiversity (Pyšek 1998; Aronson et al. 2014; Ceplová et al. 2017). Similar relationships have also been reported at finer spatial scales within cities (Crowe 1979; Matthies et al. 2015; Figueroa et al. 2018; Wang et al. 2021). Nevertheless, our analyses indicate that demographic and economic dimensions of city size (population size and GDP) produced the strongest support for the interaction predicted by the Urban Island Biogeography Framework. Population size and GDP likely capture processes that extend beyond habitat availability. Both variables reflect the intensity of human movement, trade and economic exchange that connect cities through transportation networks (Banks et al. 2015; Olden et al. 2021). Such connectivity increases propagule pressure, enhances opportunities for long-distance dispersal and promotes repeated introduction events (Banks et al. 2015; Cubino et al. 2015; O’Malia et al. 2018). At the same time, larger and wealthier cities typically contain greater environmental heterogeneity and more ornamental plantings, increasing establishment opportunities for alien species (Hope et al. 2003; Hui et al. 2017; Chamberlain et al. 2020; Yücedag & Asik 2023). Although these mechanisms were not measured directly, they provide plausible biological explanations for why socio-economic measures consistently outperformed purely spatial measures of urban extent.

Interestingly, built-up area did not support a significant interaction with geographic isolation. This finding suggests that the amount of impervious urban infrastructure alone may be a poor surrogate for processes governing plant invasions. Unlike population size or economic activity, built-up area does not necessarily reflect propagule supply, transport intensity or habitat suitability. Consequently, not all measures of urbanization are equally informative for predicting biological invasions.

### Introduction-hub identity modifies the urban size-isolation relationship

Our analyses further demonstrate that the relationship between urban size and isolation depends on the identity of the introduction hub. Across nearly all urban size metrics, Washington, representing our North American introduction hub, was the only source region exhibiting consistently negative and significant three-way interactions, indicating that the positive influence of urban size weakened with increasing distance from this introduction hub. This finding challenges an implicit assumption of classical IBT that source pools are functionally equivalent. Instead, our results indicate that the ecological consequences of geographic isolation depend not only on recipient-city characteristics but also on the identity of the donor region.

The mechanisms underlying this pattern cannot be resolved directly from our analyses, but several explanations are plausible. Richardson et al. (2025), using the same global database, showed that North America contributes substantially more alien plant species to South America than it receives in return, suggesting perhaps a strong regional exchange within the Americas. Such regional concentration of propagule exchange may produce steeper declines in colonization success with increasing distance from the Americas. Alternatively, historical trade relationships, transportation networks, donor species pools or differences in introduction history may generate source-specific dispersal patterns. Although climatic similarity may also contribute, the absence of comparable patterns for Brussels suggests that climate alone is unlikely to explain the observed interactions. Our findings therefore suggest that introduction hubs are not interchangeable sources of colonists. Rather, each donor region appears to possess its own distance-colonization relationship shaped by its historical, biogeographic and socio-economic context (Celesti-Grapow et al. 2006, Pyšek 1998; Potgieter et al. 2024). Incorporating introduction-hub identity into the Urban Island Biogeography Framework therefore represents an important extension of classical IBT, which traditionally treats isolation solely as a geographic property of recipient islands (La Sorte et al. 2014; Yang et al. 2015; Sobrinho Soares et al. 2021).

### Implications for predicting urban biological invasions

Our findings provide empirical support for the Urban Island Biogeography Framework proposed here. By integrating urban size, geographic isolation and introduction-hub identity, the framework extends classical IBT to contemporary urban ecosystems where dispersal is increasingly governed by human activities. Rather than replacing geographic isolation, anthropogenic connectivity modifies how isolation influences species accumulation, particularly in large and economically important cities.

From an applied perspective, our results suggest that invasion risk would be greatest in cities combining large population size, strong economic activity and direct connections to major introduction hubs. Consequently, urban biosecurity strategies should prioritize highly connected cities and major transportation gateways rather than relying solely on geographic proximity to donor regions. As global transportation networks continue to expand, incorporating anthropogenic connectivity into island biogeography theory may improve predictions of future urban invasions (Hulme et al. 2026).

## Conclusions

Our analyses show that alien plant richness in cities is governed primarily by the interaction between urban size and geographic isolation, while the magnitude of this relationship depends on the identity of the introduction hub. These findings demonstrate that the core principles of Island Biogeography Theory remain applicable in urban ecosystems but require explicit incorporation of anthropogenic connectivity and source-region effects. The proposed Urban Island Biogeography Framework provides a conceptual extension of classical IBT by integrating spatial, demographic and socio-economic dimensions of cities with introduction history into a unified framework for explaining global variation in urban alien plant richness. Beyond providing a mechanistic explanation for current invasion patterns, the framework generates testable predictions for understanding and managing biological invasions in an increasingly urbanized world.

## Supporting information

Supplemental data and R script

## Acknowledgements

The African Centre for DNA Barcoding is a beneficiary of the URC funding from University of Johannesburg. The graphical abstract was drawn using AI.

## Author contributions

KY did the conceptualization and designed the study and methodology; LS collected the data; LS and KY analysed the data; LS wrote the first draft. KY contributed critically to the drafts and gave final approval.

## Data availability

Data available in Figshare (Sedibana and Yessoufou 2026, DOI: 10.6084/m9.figshare.33176510)

## SUPPLEMENTAL INFORMATION

Data analyzed and R script used are avaibale in FigShare, see Sedibana and Yessoufou 2026, DOI: 10.6084/m9.figshare.33176510)

