## Supplemental data and R script for "Island Biogeography Theory-inspired predictions reveal that urban non-native plant richness is source dependent and defies classical isolation predictions": R scripts_obj1.R

```
###############################################################################
## Topic: Data Analysis testing island biogeography theory to explain alien species richness in cities
## levels: PhD student project - University of Johhanesburg, South Africa
## Kowiyou Yessoufou |
## April 19th, 2026
###############################################################################

rm(list=ls(all=TRUE))  ## Clear all

## Set working directory
 file.choose()
#####################################################
##### PART A. Measurement data: Linear Regression *******************
#####################################################
data_obj1<-read.csv2("/Users/kowiyou/Desktop/ACDB/students/PhD students/2026/phd/Lesibana/paper 1/island biogeogr/Objective_1_dataset_f.csv",sep=",",header=T)
str(data_obj1)

# R reads some numerical as characters. Lets tell R to read as numerical
data_obj1$distance_km<- as.numeric(data_obj1$distance_km)
data_obj1$latitude_min <- as.numeric(data_obj1$latitude_min)
data_obj1$longitude_min <- as.numeric(data_obj1$longitude_min)
data_obj1$latitude_max<- as.numeric(data_obj1$latitude_max)
data_obj1$longitude_max<- as.numeric(data_obj1$longitude_max)
data_obj1$latitude_centroid<- as.numeric(data_obj1$latitude_centroid)
data_obj1$longitude_centroid<- as.numeric(data_obj1$longitude_centroid)
data_obj1$total_build_up_75<- as.numeric(data_obj1$total_build_up_75)
data_obj1$total_build_up_90<- as.numeric(data_obj1$total_build_up_90)
data_obj1$total_build_up_00<- as.numeric(data_obj1$total_build_up_00)
data_obj1$total_build_up_15<- as.numeric(data_obj1$total_build_up_15)
data_obj1$pop_75<- as.numeric(data_obj1$pop_75)
data_obj1$pop_90<- as.numeric(data_obj1$pop_90)
data_obj1$pop_00<- as.numeric(data_obj1$pop_00)
data_obj1$pop15<- as.numeric(data_obj1$pop15)
data_obj1$build_up_per_capita_75<- as.numeric(data_obj1$build_up_per_capita_75)
data_obj1$build_up_per_capita_90<- as.numeric(data_obj1$build_up_per_capita_90)
data_obj1$build_up_per_capita_00<- as.numeric(data_obj1$build_up_per_capita_00)
data_obj1$build_up_per_capita_15<- as.numeric(data_obj1$build_up_per_capita_15)
data_obj1$GDP_90<- as.numeric(data_obj1$GDP_90)
data_obj1$GDP_00<- as.numeric(data_obj1$GDP_00)
data_obj1$GDP_15<- as.numeric(data_obj1$GDP_15)


names(data_obj1)


#==========================================================================
# modelling non-native SR 
#==========================================================================

data_obj1$dist_sc <- scale(data_obj1$distance_km)
data_obj1$target_city_area_km2_sc <- scale(data_obj1$target_city_area_km2)
data_obj1$urban_area_00_sc <- scale(data_obj1$urban_area_00)
data_obj1$total_build_up_15_sc <- scale(data_obj1$total_build_up_15)
data_obj1$build_up_per_capita_15_sc <- scale(data_obj1$build_up_per_capita_15)
data_obj1$pop15_sc <- scale(data_obj1$pop15)
data_obj1$GDP_15_sc <- scale(data_obj1$GDP_15)
# data_obj1$target_SR_sc <- scale(data_obj1$target_SR)


# then model fit

library(glmmTMB)

# city size as surface area

mod1a <- glmmTMB(
  target_SR ~ target_city_area_km2_sc * dist_sc * factor(source_city)+ (1 | country),
  family = nbinom2(),
  data = data_obj1
)

summary(mod1a)

#                                                                 Estimate Std. Error z value Pr(>|z|)    
# (Intercept)                                                   7.3765303  0.0363890  202.71   <2e-16 ***
#  target_city_area_km2_sc                                      -0.0007256  0.0330591   -0.02   0.9825    
#dist_sc                                                        0.0258024  0.0183696    1.40   0.1601    
#factor(source_city)Brussels                                    0.0152963  0.0256764    0.60   0.5514    
#factor(source_city)New_Delhi                                   0.0083285  0.0229821    0.36   0.7171    
#factor(source_city)Washington                                  0.0097519  0.0232607    0.42   0.6750    
#target_city_area_km2_sc:dist_sc                                0.0608978  0.0343220    1.77   0.0760 .  
#target_city_area_km2_sc:factor(source_city)Brussels            0.0429712  0.0367083    1.17   0.2418    
#target_city_area_km2_sc:factor(source_city)New_Delhi          -0.0143365  0.0473302   -0.30   0.7620    
#target_city_area_km2_sc:factor(source_city)Washington          0.0084968  0.0448269    0.19   0.8497    
#dist_sc:factor(source_city)Brussels                           -0.0223157  0.0251607   -0.89   0.3751    
#dist_sc:factor(source_city)New_Delhi                          -0.0045011  0.0223119   -0.20   0.8401    
#dist_sc:factor(source_city)Washington                         -0.0462251  0.0310064   -1.49   0.1360    
#target_city_area_km2_sc:dist_sc:factor(source_city)Brussels   -0.0106916  0.0450547   -0.24   0.8124    
#target_city_area_km2_sc:dist_sc:factor(source_city)New_Delhi  -0.0104734  0.0413792   -0.25   0.8002    
#target_city_area_km2_sc:dist_sc:factor(source_city)Washington -0.1062355  0.0461974   -2.30   0.0215 *   
  
  

# city size as urban area

mod1b <- glmmTMB(
  target_SR ~ urban_area_00_sc * dist_sc * factor(source_city) + (1 | country),
  family = nbinom2(),
  data = data_obj1
)

summary(mod1b)

#                                                       Estimate Std. Error z value Pr(>|z|)    
# (Intercept)                                            7.376338   0.036388  202.71   <2e-16 ***
#  urban_area_00_sc                                     -0.004839   0.033029   -0.15   0.8835    
#dist_sc                                                 0.025970   0.018304    1.42   0.1560    
#factor(source_city)Brussels                             0.015035   0.025634    0.59   0.5575    
#factor(source_city)New_Delhi                            0.008215   0.022916    0.36   0.7200    
#factor(source_city)Washington                           0.009884   0.023200    0.43   0.6701    
#urban_area_00_sc:dist_sc                                0.064703   0.034410    1.88   0.0601 .  
#urban_area_00_sc:factor(source_city)Brussels            0.044380   0.036783    1.21   0.2276    
#urban_area_00_sc:factor(source_city)New_Delhi          -0.016199   0.048126   -0.34   0.7364    
#urban_area_00_sc:factor(source_city)Washington          0.010359   0.045750    0.23   0.8209    
#dist_sc:factor(source_city)Brussels                    -0.022707   0.025073   -0.91   0.3651    
#dist_sc:factor(source_city)New_Delhi                   -0.004650   0.022245   -0.21   0.8344    
#dist_sc:factor(source_city)Washington                  -0.046721   0.030913   -1.51   0.1307    
#urban_area_00_sc:dist_sc:factor(source_city)Brussels   -0.010564   0.045578   -0.23   0.8167    
#urban_area_00_sc:dist_sc:factor(source_city)New_Delhi  -0.010908   0.041850   -0.26   0.7944    
#urban_area_00_sc:dist_sc:factor(source_city)Washington -0.111412   0.047213   -2.36   0.0183 *   

# city size as number of urban centres

mod1c <- glmmTMB(
  target_SR ~ urban_centre_00 * dist_sc * factor(source_city) + (1 | country),
  family = nbinom2(),
  data = data_obj1
)

summary(mod1c)

#                                                       Estimate Std. Error z value Pr(>|z|)    
# (Intercept)                                            7.420158   0.052933  140.18   <2e-16 ***
#  urban_centre_00                                       -0.048217   0.044132   -1.09   0.2746    
#dist_sc                                               -0.042724   0.035020   -1.22   0.2225    
#factor(source_city)Brussels                           -0.052047   0.054426   -0.96   0.3389    
#factor(source_city)New_Delhi                          -0.020554   0.052082   -0.39   0.6931    
#factor(source_city)Washington                         -0.032472   0.051527   -0.63   0.5286    
#urban_centre_00:dist_sc                                0.076971   0.037153    2.07   0.0383 *  
#  urban_centre_00:factor(source_city)Brussels            0.078709   0.056785    1.39   0.1657    
#urban_centre_00:factor(source_city)New_Delhi           0.035603   0.054348    0.66   0.5124    
#urban_centre_00:factor(source_city)Washington          0.050109   0.053645    0.93   0.3503    
#dist_sc:factor(source_city)Brussels                    0.011447   0.050041    0.23   0.8191    
#dist_sc:factor(source_city)New_Delhi                   0.006273   0.045268    0.14   0.8898    
#dist_sc:factor(source_city)Washington                  0.059401   0.054293    1.09   0.2739    
#urban_centre_00:dist_sc:factor(source_city)Brussels   -0.035589   0.052575   -0.68   0.4985    
#urban_centre_00:dist_sc:factor(source_city)New_Delhi  -0.006219   0.049070   -0.13   0.8991    
#urban_centre_00:dist_sc:factor(source_city)Washington -0.116664   0.053622   -2.18   0.0296 *
  --- 


# city size as total build_up 2015

mod1d <- glmmTMB(
  target_SR ~ total_build_up_15_sc * dist_sc * factor(source_city) + (1 | country),
  family = nbinom2(),
  data = data_obj1
)

summary(mod1d)

#                                                             Estimate Std. Error z value Pr(>|z|)    
#(Intercept)                                                 7.3763565  0.0364569  202.33   <2e-16 ***
#  total_build_up_15_sc                                       -0.0009391  0.0375774   -0.02   0.9801    
#dist_sc                                                     0.0251483  0.0184025    1.37   0.1718    
#factor(source_city)Brussels                                 0.0155394  0.0257338    0.60   0.5459    
#factor(source_city)New_Delhi                                0.0084948  0.0231926    0.37   0.7142    
#factor(source_city)Washington                               0.0105981  0.0235603    0.45   0.6528    
#total_build_up_15_sc:dist_sc                                0.0621226  0.0393237    1.58   0.1142    
#total_build_up_15_sc:factor(source_city)Brussels            0.0417836  0.0419353    1.00   0.3191    
#total_build_up_15_sc:factor(source_city)New_Delhi          -0.0139447  0.0581934   -0.24   0.8106    
#total_build_up_15_sc:factor(source_city)Washington          0.0131205  0.0546032    0.24   0.8101    
#dist_sc:factor(source_city)Brussels                        -0.0214851  0.0252187   -0.85   0.3942    
#dist_sc:factor(source_city)New_Delhi                       -0.0037621  0.0223985   -0.17   0.8666    
#dist_sc:factor(source_city)Washington                      -0.0459387  0.0310082   -1.48   0.1385    
#total_build_up_15_sc:dist_sc:factor(source_city)Brussels   -0.0117699  0.0541048   -0.22   0.8278    
#total_build_up_15_sc:dist_sc:factor(source_city)New_Delhi  -0.0131716  0.0493676   -0.27   0.7896    
#total_build_up_15_sc:dist_sc:factor(source_city)Washington -0.1034954  0.0565707   -1.83   0.0673 .  


# city size as population

mod1e <- glmmTMB(
  target_SR ~ pop15_sc * dist_sc * factor(source_city) + (1 | country),
  family = nbinom2(),
  data = data_obj1
)

summary(mod1e)

#                                                 Estimate Std. Error z value Pr(>|z|)    
# (Intercept)                                     7.3760876  0.0362887  203.26  < 2e-16 ***
#  pop15_sc                                        0.0009529  0.0220182    0.04 0.965479    
#dist_sc                                         0.0254124  0.0180901    1.40 0.160089    
#factor(source_city)Brussels                     0.0161244  0.0255118    0.63 0.527363    
#factor(source_city)New_Delhi                    0.0093616  0.0226518    0.41 0.679400    
#factor(source_city)Washington                   0.0090140  0.0228194    0.40 0.692831    
#pop15_sc:dist_sc                                0.0523086  0.0236158    2.21 0.026761 *  
#  pop15_sc:factor(source_city)Brussels            0.0416404  0.0262869    1.58 0.113176    
#pop15_sc:factor(source_city)New_Delhi          -0.0138510  0.0295646   -0.47 0.639428    
#pop15_sc:factor(source_city)Washington         -0.0105873  0.0300877   -0.35 0.724928    
#dist_sc:factor(source_city)Brussels            -0.0216707  0.0248134   -0.87 0.382474    
#dist_sc:factor(source_city)New_Delhi           -0.0037251  0.0220098   -0.17 0.865601    
#dist_sc:factor(source_city)Washington          -0.0494713  0.0306126   -1.62 0.106085    
#pop15_sc:dist_sc:factor(source_city)Brussels    0.0042270  0.0287957    0.15 0.883296    
#pop15_sc:dist_sc:factor(source_city)New_Delhi  -0.0001007  0.0280663    0.00 0.997138    
#pop15_sc:dist_sc:factor(source_city)Washington -0.1199452  0.0329732   -3.64 0.000275 ***
  
  
# city size as GDP

mod1f <- glmmTMB(
  target_SR ~ GDP_15_sc * dist_sc * factor(source_city) + (1 | country),
  family = nbinom2(),
  data = data_obj1
)

summary(mod1f)

#                                                  Estimate Std. Error z value Pr(>|z|)    
# (Intercept)                                      7.3750028  0.0364332  202.43  < 2e-16 ***
#  GDP_15_sc                                       -0.0199689  0.0362206   -0.55  0.58142    
#dist_sc                                          0.0267765  0.0182582    1.47  0.14250    
#factor(source_city)Brussels                      0.0167413  0.0255863    0.65  0.51291    
#factor(source_city)New_Delhi                     0.0079790  0.0228927    0.35  0.72744    
#factor(source_city)Washington                    0.0082142  0.0233643    0.35  0.72516    
#GDP_15_sc:dist_sc                                0.0829680  0.0387796    2.14  0.03240 *  
#  GDP_15_sc:factor(source_city)Brussels            0.0573818  0.0398202    1.44  0.14958    
#GDP_15_sc:factor(source_city)New_Delhi          -0.0211171  0.0531818   -0.40  0.69131    
#GDP_15_sc:factor(source_city)Washington          0.0007565  0.0556081    0.01  0.98915    
#dist_sc:factor(source_city)Brussels             -0.0230278  0.0249413   -0.92  0.35586    
#dist_sc:factor(source_city)New_Delhi            -0.0043518  0.0222087   -0.20  0.84465    
#dist_sc:factor(source_city)Washington           -0.0495094  0.0308046   -1.61  0.10801    
#GDP_15_sc:dist_sc:factor(source_city)Brussels   -0.0212169  0.0477472   -0.44  0.65678    
#GDP_15_sc:dist_sc:factor(source_city)New_Delhi  -0.0151421  0.0469169   -0.32  0.74689    
#GDP_15_sc:dist_sc:factor(source_city)Washington -0.1568372  0.0586471   -2.67  0.00749 **


# ===============================================
# FIGURES OF ALL SIGNIFICANT RELATIONSHIPS
# ===============================================

###############################################################
# Alien richness ~ Distance × City size × Source city
###############################################################

library(glmmTMB)
library(ggeffects)
library(ggplot2)
library(dplyr)
library(patchwork)

###############################################################
# PREPARE DATA
###############################################################

data_obj1$source_city <- factor(data_obj1$source_city)

###############################################################
# IMPORTANT
# RESPONSE SHOULD BE RAW SPECIES RICHNESS
###############################################################

# Example:
# target_SR = actual alien species richness
# dist_sc = scaled distance
# pop15_sc = scaled population
# GDP_15_sc = scaled GDP

###############################################################
# FIT MODELS
###############################################################

mod_pop <- glmmTMB(
  target_SR ~ pop15_sc * dist_sc * source_city +
    (1|country),
  family = nbinom2(),
  data = data_obj1
)

mod_gdp <- glmmTMB(
  target_SR ~ GDP_15_sc * dist_sc * source_city +
    (1|country),
  family = nbinom2(),
  data = data_obj1
)

###############################################################
# PARAMETERS NEEDED TO CONVERT BACK TO KM
###############################################################

dist_mean <- mean(data_obj1$distance_km, na.rm = TRUE)
dist_sd   <- sd(data_obj1$distance_km, na.rm = TRUE)

###############################################################
# LOW / MEDIUM / HIGH POPULATION
# USING REAL DATA (10th, 50th, 90th percentiles)
###############################################################

pop_vals <- quantile(
  data_obj1$pop15_sc,
  probs = c(0.10,0.50,0.90),
  na.rm = TRUE
)

###############################################################
# LOW / MEDIUM / HIGH GDP
###############################################################

gdp_vals <- quantile(
  data_obj1$GDP_15_sc,
  probs = c(0.10,0.50,0.90),
  na.rm = TRUE
)

###############################################################
# POPULATION × DISTANCE
###############################################################

pred_pop <- ggpredict(
  mod_pop,
  terms = c(
    "dist_sc",
    "pop15_sc [10%,50%,90%]"
  ),
  bias_correction = TRUE
)
###############################################################
# CONVERT DISTANCE BACK TO KM
###############################################################

pred_pop$distance_km <-
  pred_pop$x * dist_sd + dist_mean

###############################################################
# LABELS
###############################################################

pred_pop$group <- factor(
  pred_pop$group,
  labels = c(
    "Low population (10th percentile)",
    "Medium population (50th percentile)",
    "High population (90th percentile)"
  )
)

###############################################################
# FIGURE 1
###############################################################

Fig1 <- ggplot(
  pred_pop,
  aes(
    distance_km,
    predicted,
    colour = group,
    fill = group
  )
) +
  geom_line(linewidth = 1.2) +
  geom_ribbon(
    aes(
      ymin = conf.low,
      ymax = conf.high
    ),
    alpha = 0.15,
    colour = NA
  ) +
  labs(
    x = "Distance from source city (km)",
    y = "Predicted alien plant richness",
    colour = "",
    fill = "",
    title = "A) Population moderates the effect of distance"
  ) +
  theme_bw()

###############################################################
# GDP × DISTANCE
###############################################################

pred_gdp <- ggpredict(
  mod_gdp,
  terms = c(
    "dist_sc",
    "GDP_15_sc [10%,50%,90%]"
  ),
  bias_correction = TRUE
)


pred_gdp$distance_km <-
  pred_gdp$x * dist_sd + dist_mean

pred_gdp$group <- factor(
  pred_gdp$group,
  labels = c(
    "Low GDP (10th percentile)",
    "Medium GDP (50th percentile)",
    "High GDP (90th percentile)"
  )
)

###############################################################
# FIGURE 2
###############################################################

Fig2 <- ggplot(
  pred_gdp,
  aes(
    distance_km,
    predicted,
    colour = group,
    fill = group
  )
) +
  geom_line(linewidth = 1.2) +
  geom_ribbon(
    aes(
      ymin = conf.low,
      ymax = conf.high
    ),
    alpha = 0.15,
    colour = NA
  ) +
  labs(
    x = "Distance from source city (km)",
    y = "Predicted alien plant richness",
    colour = "",
    fill = "",
    title = "B) GDP moderates the effect of distance"
  ) +
  theme_bw()

###############################################################
# POPULATION × DISTANCE × SOURCE CITY
###############################################################

pred_pop3 <- ggpredict(
  mod_pop,
  terms = c(
    "dist_sc",
    "pop15_sc [10%,50%,90%]",
    "source_city"
  ),
  bias_correction = TRUE
  
)


pred_pop3$group <- factor(
  pred_pop3$group,
  labels = c(
    "Small (10th percentile)",
    "Medium (50th percentile)",
    "High (90th percentile)"
  )
)

pred_pop3$distance_km <-
  pred_pop3$x * dist_sd + dist_mean

Fig8 <- ggplot(
  pred_pop3,
  aes(
    distance_km,
    predicted,
    colour = group
  )
) +
  geom_line(linewidth = 1.2) +
  facet_wrap(~facet, ncol = 2) +
  labs(
    x = "Distance from source city (km)",
    y = "Predicted alien plant richness",
    colour = "Population level",
    title = "D) Population × Distance × Source City"
  ) +
  theme_bw()

###############################################################
# GDP × DISTANCE × SOURCE CITY
###############################################################

pred_gdp3 <- ggpredict(
  mod_gdp,
  terms = c(
    "dist_sc",
    "GDP_15_sc [10%,50%,90%]",
    "source_city"
  ),
  bias_correction = TRUE
  
)

pred_gdp3$group <- factor(
  pred_gdp3$group,
  labels = c(
    "Small (10th percentile)",
    "Medium (50th percentile)",
    "High (90th percentile)"
  )
)

pred_gdp3$distance_km <-
  pred_gdp3$x * dist_sd + dist_mean

Fig9 <- ggplot(
  pred_gdp3,
  aes(
    distance_km,
    predicted,
    colour = group
  )
) +
  geom_line(linewidth = 1.2) +
  facet_wrap(~facet, ncol = 2) +
  labs(
    x = "Distance from source city (km)",
    y = "Predicted alien plant richness",
    colour = "GDP level",
    title = "E) GDP × Distance × Source City"
  ) +
  theme_bw()


###############################################################
# CITY SURFACE AREA × DISTANCE × SOURCE CITY
###############################################################
mod_area    <- mod1a
mod_urban   <- mod1b
mod_centres <- mod1c
mod_pop     <- mod1e
mod_gdp     <- mod1f

data_obj1$source_city <- factor(data_obj1$source_city)
mod_area <- glmmTMB(
  target_SR ~ target_city_area_km2_sc *
    dist_sc *
    source_city +
    (1|country),
  family = nbinom2(),
  data = data_obj1
)

pred_area3 <- ggpredict(
  mod_area,
  terms = c(
    "dist_sc",
    "target_city_area_km2_sc [10%,50%,90%]",
    "source_city"
  )
)


pred_area3$group <- factor(
  pred_area3$group,
  labels = c(
    "Small (10th percentile)",
    "Medium (50th percentile)",
    "Large (90th percentile)"
  )
)

pred_area3$distance_km <-
  pred_area3$x * dist_sd + dist_mean

Fig5 <- ggplot(
  pred_area3,
  aes(
    distance_km,
    predicted,
    colour = group
  )
) +
  geom_line(linewidth = 1.2) +
  facet_wrap(~facet, ncol = 2) +
  labs(
    x = "Distance from source city (km)",
    y = "Predicted alien plant richness",
    colour = "City surface area",
    title = "A) City surface area × Distance × Source City"
  ) +
  theme_bw()


###############################################################
# URBAN AREA × DISTANCE × SOURCE CITY
###############################################################

mod_urban <- glmmTMB(
  target_SR ~ urban_area_00_sc *
    dist_sc *
    source_city +
    (1|country),
  family = nbinom2(),
  data = data_obj1)
  
pred_urban3 <- ggpredict(
  mod_urban,
  terms = c(
    "dist_sc",
    "urban_area_00_sc [10%,50%,90%]",
    "source_city"
  ),
  bias_correction = TRUE
)

pred_urban3$group <- factor(
  pred_urban3$group,
  labels = c(
    "Small (10th percentile)",
    "Medium (50th percentile)",
    "Large (90th percentile)"
  )
)

pred_urban3$distance_km <-
  pred_urban3$x * dist_sd + dist_mean

Fig6 <- ggplot(
  pred_urban3,
  aes(
    distance_km,
    predicted,
    colour = group
  )
) +
  geom_line(linewidth = 1.2) +
  facet_wrap(~facet, ncol = 2) +
  labs(
    x = "Distance from source city (km)",
    y = "Predicted alien plant richness",
    colour = "Urban area",
    title = "B) Urban area × Distance × Source City"
  ) +
  theme_bw()


###############################################################
# NUMBER OF URBAN CENTRES × DISTANCE × SOURCE CITY
###############################################################

mod_centres <- glmmTMB(
  target_SR ~ urban_centre_00 *
    dist_sc *
    source_city +
    (1|country),
  family = nbinom2(),
  data = data_obj1)

pred_centres3 <- ggpredict(
  mod_centres,
  terms = c(
    "dist_sc",
    "urban_centre_00 [10%,50%,90%]",
    "source_city"
  ),
  bias_correction = TRUE
)

pred_centres3$group <- factor(
  pred_centres3$group,
  labels = c(
    "Few (10th percentile)",
    "Intermediate (50th percentile)",
    "Many (90th percentile)"
  )
)

pred_centres3$distance_km <-
  pred_centres3$x * dist_sd + dist_mean

Fig7 <- ggplot(
  pred_centres3,
  aes(
    distance_km,
    predicted,
    colour = group
  )
) +
  geom_line(linewidth = 1.2) +
  facet_wrap(~facet, ncol = 2) +
  labs(
    x = "Distance from source city (km)",
    y = "Predicted alien plant richness",
    colour = "Urban centres",
    title = "C) Number of urban centres × Distance × Source City"
  ) +
  theme_bw()


###############################################################
# DISPLAY
###############################################################

#gridExtra::grid.arrange(Fig1,Fig2,ncol=1)

gridExtra::grid.arrange(Fig5,Fig6,Fig7,ncol=3)


gridExtra::grid.arrange(Fig8,Fig9, ncol=2)

### summarizes all interaction terms as forest plot


library(tidyverse)
library(ggplot2)

############################################################
## NEGATIVE BINOMIAL GLMM EFFECTS
############################################################

forest <- tribble(
  ~CitySize,~Effect,~Estimate,~SE,~P,
  
  #----------------------------------------------------------
  # Surface area
  #----------------------------------------------------------
  "Surface area","City size",-0.0007256,0.0330591,0.9825,
  "Surface area","Isolation",0.0258024,0.0183696,0.1601,
  "Surface area","IBT interaction",0.0608978,0.0343220,0.0760,
  "Surface area","Source-city interaction",-0.1062355,0.0461974,0.0215,
  
  #----------------------------------------------------------
  # Urban area
  #----------------------------------------------------------
  "Urban area","City size",-0.0048390,0.0330290,0.8835,
  "Urban area","Isolation",0.0259700,0.0183040,0.1560,
  "Urban area","IBT interaction",0.0647030,0.0344100,0.0601,
  "Urban area","Source-city interaction",-0.1114120,0.0472130,0.0183,
  
  #----------------------------------------------------------
  # Urban centres
  #----------------------------------------------------------
  "Urban centres","City size",-0.0482170,0.0441320,0.2746,
  "Urban centres","Isolation",-0.0427240,0.0350200,0.2225,
  "Urban centres","IBT interaction",0.0769710,0.0371530,0.0383,
  "Urban centres","Source-city interaction",-0.1166640,0.0536220,0.0296,
  
  #----------------------------------------------------------
  # Built-up area
  #----------------------------------------------------------
  "Built-up area","City size",-0.0009391,0.0375774,0.9801,
  "Built-up area","Isolation",0.0251483,0.0184025,0.1718,
  "Built-up area","IBT interaction",0.0621226,0.0393237,0.1142,
  "Built-up area","Source-city interaction",-0.1034954,0.0565707,0.0673,
  
  #----------------------------------------------------------
  # Population
  #----------------------------------------------------------
  "Population","City size",0.0009529,0.0220182,0.9655,
  "Population","Isolation",0.0254124,0.0180901,0.1601,
  "Population","IBT interaction",0.0523086,0.0236158,0.0268,
  "Population","Source-city interaction",-0.1199452,0.0329732,0.000275,
  
  #----------------------------------------------------------
  # GDP
  #----------------------------------------------------------
  "GDP","City size",-0.0199689,0.0362206,0.5814,
  "GDP","Isolation",0.0267765,0.0182582,0.1425,
  "GDP","IBT interaction",0.0829680,0.0387796,0.0324,
  "GDP","Source-city interaction",-0.1568372,0.0586471,0.00749
)

############################################################
## 95% CONFIDENCE INTERVALS
############################################################

forest <- forest %>%
  mutate(
    Lower = Estimate - 1.96*SE,
    Upper = Estimate + 1.96*SE,
    sig = case_when(
      P < 0.001 ~ "***",
      P < 0.01  ~ "**",
      P < 0.05  ~ "*",
      P < 0.10  ~ "†",
      TRUE ~ ""
    )
  )

############################################################
## ORDER
############################################################

forest$Effect <- factor(
  forest$Effect,
  levels=c(
    "City size",
    "Isolation",
    "IBT interaction",
    "Source-city interaction"
  )
)

forest$CitySize <- factor(
  forest$CitySize,
  levels=c(
    "Surface area",
    "Urban area",
    "Urban centres",
    "Built-up area",
    "Population",
    "GDP"
  )
)

############################################################
## FOREST PLOT
############################################################

ggplot(forest,
       aes(x=Estimate,
           y=CitySize,
           colour=Effect))+
  
  geom_vline(
    xintercept=0,
    linetype=2,
    colour="grey50"
  )+
  
  geom_errorbarh(
    aes(xmin=Lower,
        xmax=Upper),
    height=0.20,
    linewidth=0.8,
    position=position_dodge(width=0.7)
  )+
  
  geom_point(
    size=3.5,
    position=position_dodge(width=0.7)
  )+
  
  geom_text(
    aes(label=sig),
    nudge_x=0.018,
    size=5,
    show.legend=FALSE,
    position=position_dodge(width=0.7)
  )+
  
  scale_colour_manual(values=c(
    "City size"="grey30",
    "Isolation"="#2C7BB6",
    "IBT interaction"="#FDAE61",
    "Source-city interaction"="#D7191C"
  ))+
  
  labs(
    x=expression(beta~coefficient~"(95% CI)"),
    y=NULL,
    colour=NULL
  )+
  
  theme_bw(base_size=15)+
  
  theme(
    legend.position="bottom",
    panel.grid.minor=element_blank(),
    panel.grid.major.y=element_blank(),
    legend.text=element_text(size=12),
    axis.text.y=element_text(face="bold")
  )


########################
# by how much alien SR would increase if city size and distance increase simultaneously by 1 unit?

## CONDITIONAL MARGINAL EFFECTS OF URBAN SIZE
## ACROSS THE OBSERVED DISTANCE GRADIENT
###############################################################

library(marginaleffects)
library(dplyr)
library(ggplot2)

###############################################################
## Mean and SD used to standardize distance
###############################################################

dist_mean <- mean(data_obj1$distance_km, na.rm = TRUE)
dist_sd   <- sd(data_obj1$distance_km, na.rm = TRUE)

###############################################################
## Function
###############################################################

conditional_effect <- function(model,
                               size_var,
                               metric_name,
                               data){
  
  #############################################################
  ## Distance sequence (standardized)
  #############################################################
  
  d.seq <- seq(
    min(data$dist_sc, na.rm = TRUE),
    max(data$dist_sc, na.rm = TRUE),
    length.out = 100
  )
  
  #############################################################
  ## Prediction grid
  #############################################################
  
  grid <- datagrid(
    model = model,
    dist_sc = d.seq,
    source_city = levels(data$source_city)
  )
  
  #############################################################
  ## Marginal effects
  #############################################################
  
  out <- slopes(
    model,
    variables = size_var,
    newdata = grid,
    type = "response"
  )
  
  out <- as.data.frame(out)
  
  #############################################################
  ## Convert standardized distance back to km
  #############################################################
  
  out$Distance_km <- out$dist_sc * dist_sd + dist_mean
  
  out$UrbanMetric <- metric_name
  
  out
  
}


### table showing by how much alien increases with a simultaneous increase in distance and city size by a unit


###############################################################
## PREDICTED CHANGE IN ALIEN RICHNESS FOR MEANINGFUL
## RAW INCREMENTS OF URBAN SIZE AND DISTANCE
###############################################################

library(dplyr)

predict_change_raw <- function(model,
                               size_var_sc,
                               size_var_raw,
                               raw_increment,
                               metric_name,
                               data,
                               standardized = TRUE){
  
  #############################################################
  ## Convert raw increment to standardized units
  #############################################################
  
  if(standardized){
    
    delta_size <-
      raw_increment /
      sd(data[[size_var_raw]], na.rm = TRUE)
    
  } else{
    
    ## variable already in raw units
    delta_size <- raw_increment
    
  }
  
  ## distance increment = 1000 km
  
  delta_dist <-
    1000 /
    sd(data$distance_km, na.rm = TRUE)
  
  #############################################################
  ## Predictions
  #############################################################
  
  hubs <- levels(data$source_city)
  
  Results <- lapply(hubs, function(h){
    
    new0 <- data[1,]
    
    ## numeric predictors at their mean
    num <- sapply(new0,is.numeric)
    
    for(v in names(new0)[num])
      new0[[v]] <- mean(data[[v]], na.rm=TRUE)
    
    ###########################################################
    ## reference values
    ###########################################################
    
    new0[[size_var_sc]] <- mean(data[[size_var_sc]], na.rm=TRUE)
    
    new0$dist_sc <- mean(data$dist_sc, na.rm=TRUE)
    
    new0$source_city <- h
    
    ###########################################################
    ## increased values
    ###########################################################
    
    new1 <- new0
    
    new1[[size_var_sc]] <-
      new0[[size_var_sc]] + delta_size
    
    new1$dist_sc <-
      new0$dist_sc + delta_dist
    
    ###########################################################
    ## predictions
    ###########################################################
    
    SR0 <- predict(
      model,
      newdata=new0,
      type="response",
      re.form=NA
    )
    
    SR1 <- predict(
      model,
      newdata=new1,
      type="response",
      re.form=NA
    )
    
    data.frame(
      
      Metric = metric_name,
      
      SourceCity = h,
      
      SizeIncrease = raw_increment,
      
      DistanceIncrease = 1000,
      
      Reference = SR0,
      
      Predicted = SR1,
      
      Increase = SR1-SR0
      
    )
    
  })
  
  bind_rows(Results)
  
}

###############################################################
## Run all models
###############################################################

Table_raw <- bind_rows(
  
  predict_change_raw(
    mod1a,
    "target_city_area_km2_sc",
    "target_city_area_km2",
    100,
    "City surface area",
    data_obj1
  ),
  
  predict_change_raw(
    mod1b,
    "urban_area_00_sc",
    "urban_area_00",
    100,
    "Urban area",
    data_obj1
  ),
  
  predict_change_raw(
    mod1c,
    "urban_centre_00",
    "urban_centre_00",
    1,
    "Urban centres",
    data_obj1,
    standardized = FALSE
  ),
  
  predict_change_raw(
    mod1d,
    "total_build_up_15_sc",
    "total_build_up_15",
    100,
    "Built-up area",
    data_obj1
  ),
  
  predict_change_raw(
    mod1e,
    "pop15_sc",
    "pop15",
    1000000,
    "Population",
    data_obj1
  ),
  
  predict_change_raw(
    mod1f,
    "GDP_15_sc",
    "GDP_15",
    10000000000,
    "GDP",
    data_obj1
  )
  
)

Table_raw

#               Metric SourceCity SizeIncrease DistanceIncrease Reference Predicted   Increase
#1  City surface area    Beijing        1e+02             1000  1598.035  1616.689  18.653231
#2  City surface area   Brussels        1e+02             1000  1622.667  1652.858  30.190287
#3  City surface area  New_Delhi        1e+02             1000  1611.400  1619.514   8.113565
#4  City surface area Washington        1e+02             1000  1613.695  1602.962 -10.732986
#5         Urban area    Beijing        1e+02             1000  1597.729  1615.270  17.541782
#6         Urban area   Brussels        1e+02             1000  1621.933  1652.998  31.065616
#7         Urban area  New_Delhi        1e+02             1000  1610.907  1616.190   5.282852
#8         Urban area Washington        1e+02             1000  1613.600  1601.180 -12.419272
#9      Urban centres    Beijing        1e+00             1000  1604.303  1568.506 -35.796794
#10     Urban centres   Brussels        1e+00             1000  1624.938  1694.809  69.870630
#11     Urban centres  New_Delhi        1e+00             1000  1618.433  1637.489  19.056016
#12     Urban centres Washington        1e+00             1000  1618.481  1597.986 -20.494859
#13     Built-up area    Beijing        1e+02             1000  1597.758  1618.667  20.909363
#14     Built-up area   Brussels        1e+02             1000  1622.780  1661.274  38.494631
#15     Built-up area  New_Delhi        1e+02             1000  1611.388  1619.104   7.715823
#16     Built-up area Washington        1e+02             1000  1614.781  1606.862  -7.919085
#17        Population    Beijing        1e+06             1000  1597.328  1635.935  38.607404
#18        Population   Brussels        1e+06             1000  1623.293  1737.945 114.652477
#19        Population  New_Delhi        1e+06             1000  1612.352  1623.117  10.765463
#20        Population Washington        1e+06             1000  1611.792  1551.061 -60.730610
#21               GDP    Beijing        1e+10             1000  1595.596  1607.910  12.314195
#22               GDP   Brussels        1e+10             1000  1622.533  1655.583  33.049668
#23               GDP  New_Delhi        1e+10             1000  1608.379  1604.476  -3.902144
#24               GDP Washington        1e+10             1000  1608.757  1577.245 -31.511535

write.table(Table_raw,"Table2_ibt")

# graphicaall representation of the change

###############################################################
## FOREST PLOT OF PREDICTED CHANGES
###############################################################

library(ggplot2)
library(dplyr)

###############################################################
## Prepare data
###############################################################

PlotDat <- Table_raw %>%
  mutate(
    
    SourceCity = factor(
      SourceCity,
      levels = c("Beijing","Brussels","New_Delhi","Washington"),
      labels = c("Beijing","Brussels","New Delhi","Washington")
    ),
    
    Metric = factor(
      Metric,
      levels = c(
        "Population",
        "GDP",
        "Built-up area",
        "Urban area",
        "City surface area",
        "Urban centres"
      )
    ),
    
    Increment = case_when(
      Metric == "Population" ~ "+1 million inhabitants",
      Metric == "GDP" ~ "+US$10 billion",
      Metric == "Built-up area" ~ "+100 km²",
      Metric == "Urban area" ~ "+100 km²",
      Metric == "City surface area" ~ "+100 km²",
      Metric == "Urban centres" ~ "+1 urban centre"
    ),
    
    MetricLabel = paste0(Metric,"\n(",Increment,")")
  )

###############################################################
## Forest plot
###############################################################

Fig_Forest <-
  
  ggplot(
    PlotDat,
    aes(
      x = Increase,
      y = MetricLabel,
      colour = SourceCity
    )
  )+
  
  geom_vline(
    xintercept = 0,
    linetype = "dashed",
    colour = "grey40",
    linewidth = 0.8
  )+
  
  geom_point(
    size = 3.8,
    position = position_dodge(width = 0.6)
  )+
  
  ## horizontal line from zero to estimate
  geom_segment(
    aes(
      x = 0,
      xend = Increase,
      y = MetricLabel,
      yend = MetricLabel
    ),
    linewidth = 0.8,
    position = position_dodge(width = 0.6),
    alpha = 0.8
  )+
  
  scale_colour_brewer(
    palette = "Dark2"
  )+
  
  labs(
    x = "Predicted change in alien plant richness",
    y = NULL,
    colour = "Introduction hub"
  )+
  
  theme_bw(base_size = 15)+
  
  theme(
    
    legend.position = "bottom",
    
    legend.title = element_text(face = "bold"),
    
    axis.text.y = element_text(face = "bold"),
    
    panel.grid.major.y = element_blank(),
    
    panel.grid.minor = element_blank(),
    
    panel.grid.major.x = element_line(colour = "grey90")
    
  )

Fig_Forest


############################################################
# GLOBAL MAPS: Alien species richness & Area (scaled data)
############################################################


library(sf)
library(ggplot2)
library(dplyr)
library(rnaturalearth)
library(rnaturalearthdata)
library(viridis)
library(patchwork)
library(ggeffects)

###############################################################
# WORLD MAP
###############################################################

world <- ne_countries(
  scale = "medium",
  returnclass = "sf"
)

###############################################################
# SOURCE CITY COORDINATES
###############################################################

sources <- data.frame(
  source_city = factor(
    c("Beijing",
      "Brussels",
      "New_Delhi",
      "Washington"),
    levels = levels(data_obj1$source_city)
  ),
  source_lon = c(
    116.4074,
    4.3517,
    77.2090,
    -77.0369
  ),
  source_lat = c(
    39.9042,
    50.8503,
    28.6139,
    38.9072
  )
)

###############################################################
# JOIN SOURCE COORDINATES
###############################################################

links <- left_join(
  data_obj1,
  sources,
  by = "source_city"
)

###############################################################
# MODEL RESIDUALS
###############################################################

links$residuals <- NA
 
 used_rows <- complete.cases(
         data_obj1[, c(
               "target_SR",
               "pop15_sc",
               "dist_sc",
               "source_city",
               "country"
           )]
     )
 
   links$residuals[used_rows] <-
       residuals(
             mod1d, # mod1d chosed because has the lowest AIC (34329.76)
             type = "pearson"
         )
 Map4 <- ggplot() +
       geom_sf(
             data = world,
             fill = "grey95",
             colour = "grey80"
         ) +
       geom_point(
             data = subset(links, !is.na(residuals)),
             aes(
                   longitude_centroid,
                   latitude_centroid,
                   colour = residuals
               ),
             alpha = 0.8,
             size = 2
         ) +
       scale_colour_gradient2(
             low = "blue",
             mid = "white",
             high = "red",
             midpoint = 0,
             name = "Std. residual"
         ) +
   labs(
     title = "C) Spatial distribution of model residuals",
     x = NULL,
     y = NULL
   ) +
   theme_bw()
       
###############################################################
# HOTSPOTS
###############################################################

hotspots <- links %>%
  filter(
    target_SR >= quantile(
      target_SR,
      0.90,
      na.rm = TRUE
    )
  )

###############################################################
# MAP A
# GLOBAL ALIEN RICHNESS
###############################################################

Map2 <- ggplot() +
  geom_sf(
    data = world,
    fill = "grey95",
    colour = "grey80",
    linewidth = 0.2
  ) +
  geom_point(
    data = links,
    aes(
      longitude_centroid,
      latitude_centroid,
      colour = target_SR,
      size = target_SR
    ),
    alpha = 0.8
  ) +
  scale_colour_viridis_c(name = "Richness") +
  labs(
    title = "A) global distribution of urban alien plant richness",
    x = NULL,
    y = NULL
  ) +
  theme_bw()

###############################################################
# MAP D
# SOURCE-TARGET NETWORK
###############################################################

Map1 <- ggplot() +
  geom_sf(
    data = world,
    fill = "grey95",
    colour = "grey80"
  ) +
  geom_curve(
    data = links,
    aes(
      x = source_lon,
      y = source_lat,
      xend = longitude_centroid,
      yend = latitude_centroid
    ),
    alpha = 0.02,
    linewidth = 0.5,
    curvature = 0.5
  ) +
  geom_point(
    data = sources,
    aes(
      source_lon,
      source_lat
    ),
    size = 3
  ) +
  labs(
   # title = "D) Conceptual source–target network",
    x = NULL,
    y = NULL
  ) +
  theme_bw()

###############################################################
# MAP B
# HOTSPOTS
###############################################################

Map3 <- ggplot() +
  geom_sf(
    data = world,
    fill = "grey95",
    colour = "grey80"
  ) +
  geom_point(
    data = hotspots,
    aes(
      longitude_centroid,
      latitude_centroid,
      colour = target_SR,
      size = target_SR
    )
  ) +
  scale_colour_viridis_c(name = "Richness") +
  labs(
    title = "B) Global hotspots of urban alien plant richness",
    x = NULL,
    y = NULL
  ) +
  theme_bw()


###############################################################
# COMBINE ALL PANELS
###############################################################

FinalFigure <-
  (Map1 + Map2) /
  (Map3 + Map4)
  

FinalFigure +
  plot_annotation(
    title = "Global patterns of alien plant richness and the moderating effects of city size on distance-decay relationships"
  )

dev.off()
gridExtra::grid.arrange(Map3, Map4, ncol=1)

gridExtra::grid.arrange( Map1,Map2,Map3, ncol=1)

gridExtra::grid.arrange(Map2,Map3, ncol=1)

Map1
```
